# Establishment and characterization of novel autologous pair primary cultures from two Indian non-habitual tongue carcinoma patients

**DOI:** 10.1101/2022.01.25.477260

**Authors:** Nehanjali Dwivedi, G Charitha, Vijay Pillai, Moni A Kuriakose, Amritha Suresh, Manjula Das

## Abstract

Oral tongue squamous cell carcinoma (OTSCC) is one of the major causes of fatality in India owing to very high percentages of patients with smoking and chewing habits. Being highly heterogeneous in nature, every patient poses a different challenge clinically. To better understand disease progression, knowledge of cross talk between tumor stroma and the tumor cells becomes indispensable. Patient-derived in vitro cell line models are helpful to understand the complexity of diseases. However, they have very low efficiency of establishment from the tumor samples, especially the cancer associated fibroblasts (CAFs). In the present study, two novel autologous pairs have been immortalized spontaneously from non-habitual, HPV-positive patients, presented with tongue squamous cell carcinoma. The epithelial and fibroblast primary lines had typical polygonal and spindle shaped morphology, respectively. Positive staining with Pan-cytokeratin (PanCK) and Fibroblast Specific Protein (FSP-1) further confirmed their epithelial and fibroblast origin. Unique Short Tandem Repeat (STR) profile of the cultures confirmed their novelty, while the similarity of the STR profiles between the epithelial and fibroblast cells from the same patient, confirmed their autologous nature. DNA analysis revealed aneuploidy of the established cultures. Increase in the tumorigenic potential of the established epithelial cultures upon treatment with CAF-conditioned medium proved the “CAF-ness” of the established fibroblast cells. The established cultures are the first of their kind which would serve as an useful platform in understanding the cross talk between tumor-stroma and tumor, along with studying tongue cancer progression.

## Introduction

Cancer of oral cavity is ranked among the top five most frequent cancers occurring in India among both males and females. Comprising of 10.3% of all cancers, males (16.2%) are seen to be affected more by it than females (4.6%) in India (1). OTSCC distresses approximately 16,000 people a year in the US (2) and is typically related to a long history of smoking and/or heavy alcohol use (3). Even though, the smoking rates continue to drop, the incidence of squamous cell carcinoma (SCC) of the oral cavity has remained constant (4) due to the increase of OSCC patients who have never smoked or whose habit was non-significant. These individuals are often women in their mid-forties or younger (3, 5–8). HPV has been implicated in the recent rise in oropharyngeal cancers (9,10). Among the tongue cancer patients, a very small percentage has been shown to be HPV positive (11).

Knowledge of the molecular and biological characteristics of cancer cells is essential to develop therapeutic strategies or finding drug targets of cancer treatment. Particularly, the primary culture, immortalized spontaneously from patient tumors, retain the characteristics displayed by the parent tumors. Therefore, these primary cultures become a desirable source for conducting translational research in laboratory settings. The scarcity of commercial cell lines, especially CAFs, imposes the need to establish and characterise novel cell lines from patient tumors (12).

There have been various reports of establishment of oral cancer cell lines from tumors (13–15). In USA and UK initial attempts were made to establish cell lines from head and neck squamous cell carcinoma (HNSCC) patients who had undergone chemo and radiotherapy (16,17). Zhao et.al in 2011 further assembled and characterized 85 cell lines from various head and neck tumor sites (18). More recently, one primary epithelial line was established from a non-smoking patient, diagnosed with oral tongue squamous cell carcinoma (19). However, an instance wherein both the epithelial and CAF are established from the same patient are not available which will assemble a superior framework in understanding cancer progression (20).

Establishment of oral cancer cell lines from Indian patients have been attempted from various head and neck sites (13,21). In the present study, establishment of cells from tongue of two patients have been reported. The novelty of the cell lines was validated by determining their STR profiles. The epithelial cells when cultured in presence of the conditioned medium from fibroblasts, showed increased tumorigenic properties. The knowledge about the interaction of epithelial tumor cells with the static microenvironment (fibroblasts) can be deduced using such a platform. Till date, most tongue cancer cell lines have been established from patients with habits, or whose habits are unknown, supporting the need for establishing cell lines from verified non habitual patients. Thus the autologous pairs reported in the study can be used as a model system to identify theragnostic biomarkers as appropriate.

## Materials and methods

### Tumor specimen and establishment

Tumor samples were collected after obtaining prior informed consent from patients following the approval of the institutional ethics committee. Tissues were collected aseptically in RPMI-1640 (Cat # AT222A, Himedia, India) with triple strength penicillin-streptomycin (Cat# 15140122, Gibco™, U.S.A) from two 65-year-old females with no risk habits, diagnosed with squamous cell carcinoma of tongue. Patient MhCT08 had undergone neoadjuvant therapy (CT OR CTRT) and patient MhCT12 was a naive surgical sample. The clinical details of the patients are mentioned in Table I. The tissue samples were thoroughly washed 3 times with 5 minutes interval with 3X Penicillin-streptomycin followed by 10% povidone iodine solution (Win Medicare) and finally with complete growth medium. The tissue was chopped into small pieces, treated with 0.25% trypsin (Cat # 25200056, Gibco™, U.S.A) for 30 minutes at 37°C. The chopped and digested pieces were placed in a serrated 60mm petri dishes and supplemented with 10ng/μl each of human recombinant epidermal growth factor (hEGF, Cat # E9644, Sigma-Aldrich), N2 suppliment-1X (Cat # 17502048, Gibco™, U.S.A), Epilife defined growth supplement (EDGS, Cat # S0125, Gibco™, U.S.A) along with 20% FBS (Cat # RM10434, Himedia, India), in RPMI-1640 media with 1x penicillin-streptomycin. The media was changed every 48hrs to remove the dead cells, the epithelial cells were enriched by differential trypsinisation and further sub-cultured. For differential trypsinization, briefly, the cells were trypsinized for two different time points. After a minute of trypsinization, floating cells were removed and seeded in a separate flask. Since fibroblasts can detach faster than epithelial cells, this differential trypsinization technique yielded two separate cellular populations. The separated cells were grown in RPMI media with 20% FBS and no additional growth supplements. The cells were passaged for more than P50 and were characterised for cell type specificity at both early and late passages. Later passages of the cells were maintained in RPMI medium supplemented with 20% FBS and 1X penicillin-streptomycin solution.

**Table I.** Clinical and pathological details of the established cell lines. Patient MhCT08 had undergone neoadjuvant therapy (CT OR CTRT) and patient MhCT12 was a naive surgical sample.

| Cell line | Patient Age (years) | Sex | Origin of tumor | Tumor diagnosis | Oral Habit | Ethnicity |
| --- | --- | --- | --- | --- | --- | --- |
| MhCT08 | 65 | F | Tongue | Squamous cell carcinoma | None | Indian |
| MhCT12 | 65 | F | Tongue | Squamous cell carcinoma | None | Indian |

F: Female

### Hela cell culture

HeLa cells were cultured in DMEM medium (Cat # 11995-065, Gibco™, U.S.A) supplemented with 10% FBS and 1X penicillin-streptomycin solution.

### Characterization

#### Growth curve

Cells were collected and seeded at a concentration of 1×10^4^ cells per well in 6-well plates. Cell count of three random wells was taken every day up to 5 days using trypan blue (Cat # T6146, Sigma, U.S.A). The doubling time was calculated using the equation - Td = TX log2/log(N/N0), where Td is the doubling time; T is the time interval; N is the final cell number; and N0 is the initial cell number. Results show the mean ± Standard Deviation (SD) of three independent experiments.

#### Flow cytometry

Cells at a concentration of 10^6^ cells/ 100μl were washed twice in PBS, permeabilised with 0.1% triton X100 (Cat # 10655, Fisher Scientific, U.S.A) for 30 min and incubated with primary antibody (1:50) for 1 h on ice. The cell types were probed with anti-Pan cytokeratin (Cat # 4545, Cell signalling technology, U.S.A) and anti-Fibroblast specific protein (Cat # F4771, Sigma, U.S.A) as primary antibodies. Cells were then pelleted down by centrifugation at 500g for 5 minutes and washed with PBS (Cat # 10010-023, Gibco™, U.S.A) followed by incubation with the corresponding secondary antibody, anti-mouse Alexa 488 (Cat# A11029, Invitrogen, U.S.A) fluorochrome for 30 minutes at room temperature in dark conditions. For the marker expression analysis, cell sorting gates were established using unstained control population (n=1). The cells were analysed using BD FACS Canto II system.

#### DNA ploidy determination

Cells were harvested following trypsinization and resuspended in PBS containing RNase (10μg/ml) (Cat# 12091021, Invitrogen, U.S.A) and propidium iodide (40 μg/ml) (Cat # P4170, Sigma Aldrich, U.S.A) and incubated at 37°C for 30 min for DNA staining. The DNA content was compared to human mononuclear cells from peripheral blood, which served as a control for diploid human genomic DNA content. The cells were analysed using a BD FACS Canto II system to determine their fluorescence. The mean channel of cells that were in the G0 phase was divided by that of the lymphocytes to determine DNA index of each cell line. The DNA index was then used to predict ploidy of the cells.

#### Immunocytochemistry

The established cell lines were examined for the presence of HPV infection status by staining with p16 (Cat# AM540-5M, BioGenex, India) and E7 antibody (Cat# sc-58661, Santa Cruz Biotechnology, U.S.A) via immunocytochemistry. HeLa cells, a cervical cancer cell line, was used as a positive control for the experiment. Cells (5×10^3^) were grown on coverslips and fixed in 4% Paraformaldehyde for 10 minutes (Cat # GRM3660, HiMedia, India) followed by permeabilization with 0.1% triton X-100 for 10 minutes, and probed with p16 or E7 overnight. The coverslips were further processed with Dako Kit (Cat # K5007, Dako, Agilent, U.S.A). The presence of target proteins was visualised using DAB as chromogen and the cells were counter stained with Haematoxylin and mounted with DPX mountant and examined under Nikon Eclipse E200 light microscope.

#### PCR

All the samples were screened for the presence of HPV infection by PCR to amplify a 450bp fragment from the conserved region in all HPV variants with MY09/MY11 primers: 5’-CGTCCMARRGGAWACTGATC-3’ as forward and 5’-GCMCAGGGWCATAAYAATGG-3’ as the reverse primer (23). PCR was performed using 200ng of DNA along with dNTP (0.2mM each), Taq polymerase (Cat # D1806, Sigma, U.S.A), 1X reaction buffer and primers at a final concentration of 0.1μM each. Forward primer 5’-AGCCATGTACGTTGCTATCCA-3’ and reverse primer 5’-ACCGGAGTCCATCACGATG-3’ were used to amplify beta actin (amplicon 120bp). Amplifications were performed using the cycling conditions-94°C for 5 min followed by denaturation at 95°C for 1 min, annealing for 1 min (50°C for MY09/MY11 and 60°C for Beta actin), and elongation at 72°C for 1 min. A final elongation step was carried out at the end of the final cycle at 72°C for 10 minutes. DNA extracted from HeLa cell line was used as the positive control and as a negative control, no template was added to the reaction mix. The samples were subjected to agarose gel electrophoresis (1.2% gel) for amplicon visualization.

#### STR profiling

STR profiling of 10 loci was performed to establish the genomic identity, cell line identity and exclude any cross-contamination of MhCT08 and MhCT12-fibroblast and epithelial cells. STR multiplex assay was outsourced to Theracues, India and performed using GenePrint 10 (Promega). SoftGenetics GeneMarker_HID version 3.0.0 was used for analysing the results. STR data were then examined in the reference STR database of ATCC and CLASTR using the standard match threshold of 80%.

#### Estimation of proliferation

Conditioned media from CAFs was collected 48 hours after seeding. The epithelial cells were seeded at a concentration of 1×10^4^ cells per well in a 96 well plate for attachment overnight at 37°C in a humidified atmosphere with 5% CO_2_. The cells were treated with different ratios of the collected CAF conditioned medium and fresh medium, as indicated in appropriate figures, for 48 hours, and cell proliferative ability was measured using Alamar Blue Reagent (Cat # R7017, Sigma, U.S.A), prepared at a final concentration of 0.2mg/mL in sterile PBS. Per 100μL of the growth medium in a 96 well plate, 20μL of Alamar blue was added and incubated for 2 hours at 37°C. Optical Density at 570nm and 600nm were measured. Proliferation was calculated using the formula (*O2 X A1*) − (*O1 X A2*), where, O1 is the molar extinction coefficient (E) of oxidized Alamar blue at 570nm, O2 is the E of the oxidized Alamar blue at 600nm, A1 is the absorbance of test wells at 570nm and A2 is the absorbance of test wells at 600nm. From the table of oxidation coefficients of Alamar blue, values of O1 and O2 were taken as 80856 and 117216 respectively. The fold proliferation was calculated over the negative control well having media and the Alamar blue reagent without any cells. Results show the mean ± SD of three independent experiments.

#### Wound healing assay

Conditioned media from CAFs was collected for 24, 48 and 72 hours after seeding. Epithelial cells (0.3×10^6^) were seeded in each well of a 6 well plate and grown till 80% confluency. The cell monolayer was gently scraped with a sterile 200μl pipette tip and the wells were washed twice with 1X PBS to remove cell debris. The cells were then treated with different ratios of the collected conditioned medium and fresh medium, as indicated in figures. The width of the scratch was determined by images taken under a light microscope at 0 hour and 24 hours after creating the wound. The wound closure was then quantified using ImageJ software 1.53k and percent wound closure was plotted. Percentage wound healing was calculated by the formula 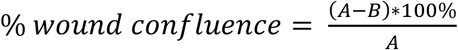, where, A is the width of the initial scratch wound and B is the width of the scratch wound at time 24h. Results show the mean ± SD of three independent experiments.

#### Invasion assay

Conditioned medium from CAFs was collected 24 and 48 hours after seeding and used as a chemo-attractant for invasion assay. Briefly, ECM gel (Cat # E1270, Sigma, U.S.A) was prepared at a final concentration of 1mg/mL in DMEM serum free media. 100μL of ECM was coated on the transwell inserts and incubated for 2 hours at 37°C to allow gel formation. 1×10^4^ cells were then seeded in serum free medium on the top chamber of the transwell insert (Cat # TCP083, HiMedia, India) with conditioned medium as the chemoattractant in the lower chamber. The cells were allowed to invade for 48 hours. For imaging, the cells on the inside of the transwell were removed gently using cotton swabs, followed by staining with crystal violet (Cat # S012, HiMedia, India) for 10 minutes. The transwell inserts were washed with PBS twice to remove any unbound crystal violet and then air-dried before imaging. For quantification of invasion, bound crystal violet was eluted by incubating the transwell inserts in 10% acetic acid (Cat # Q21057, Qualigen, U.S.A) in water with shaking for 10 minutes. The eluent was then transferred to a 96-well clear microplate, and the absorbance was read at 590nm. Results show the mean ± SD of three independent experiments.

#### Sphere formation assay

3D sphere formation was carried out as detailed by Arya et.al, 2016 (24). Briefly, 0.5 × 10^6^ cells were encapsulated in 7.5% 3D GelMA hydrogels and cultured for 14 days, with a partial medium change (normal or CAF conditioned medium) on alternative days. After 14 days, the spheroids were harvested using enzymatic degradation of the hydrogel and images were captured. Spheroid size was calculated using the formula 4/3πr^3^, where ‘r’ represents the geometric mean of the two diameters of the spheroids. Results show the mean ± SD of three independent experiments.

#### Statistical analysis

All the quantitative data is expressed as the mean ± standard error of the mean. Statistical significance was calculated using Student’s T-test. p-value less than 0.1(indicated as *) and 0.05 (indicated as **) were taken as statistically significant.

## Results

Tissue samples from both patients, coded as MhCT12 and MhCT08, were sub-cultured for more than 40 passages. Both yielded epithelial as well as cancer associated fibroblast cells and were further characterized in various ways.

### Morphology

Image analysis under light microscope revealed typical polygonal morphology of both the epithelial cells while both the fibroblast cell types were observed to have typical spindle shaped morphology as shown in Fig. 1A. The images were taken after passage 35 for each cell type.

**Figure 1.**
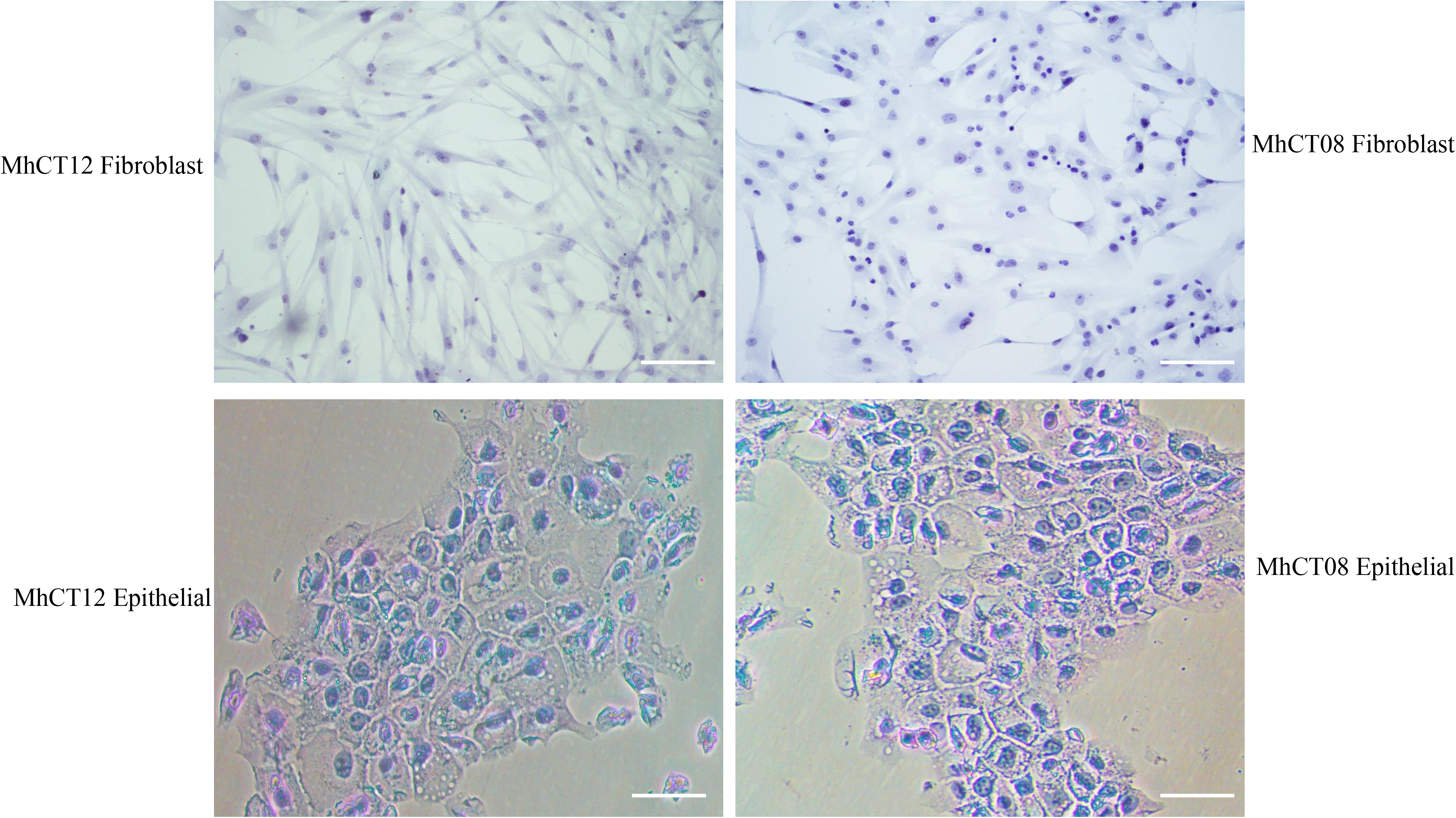
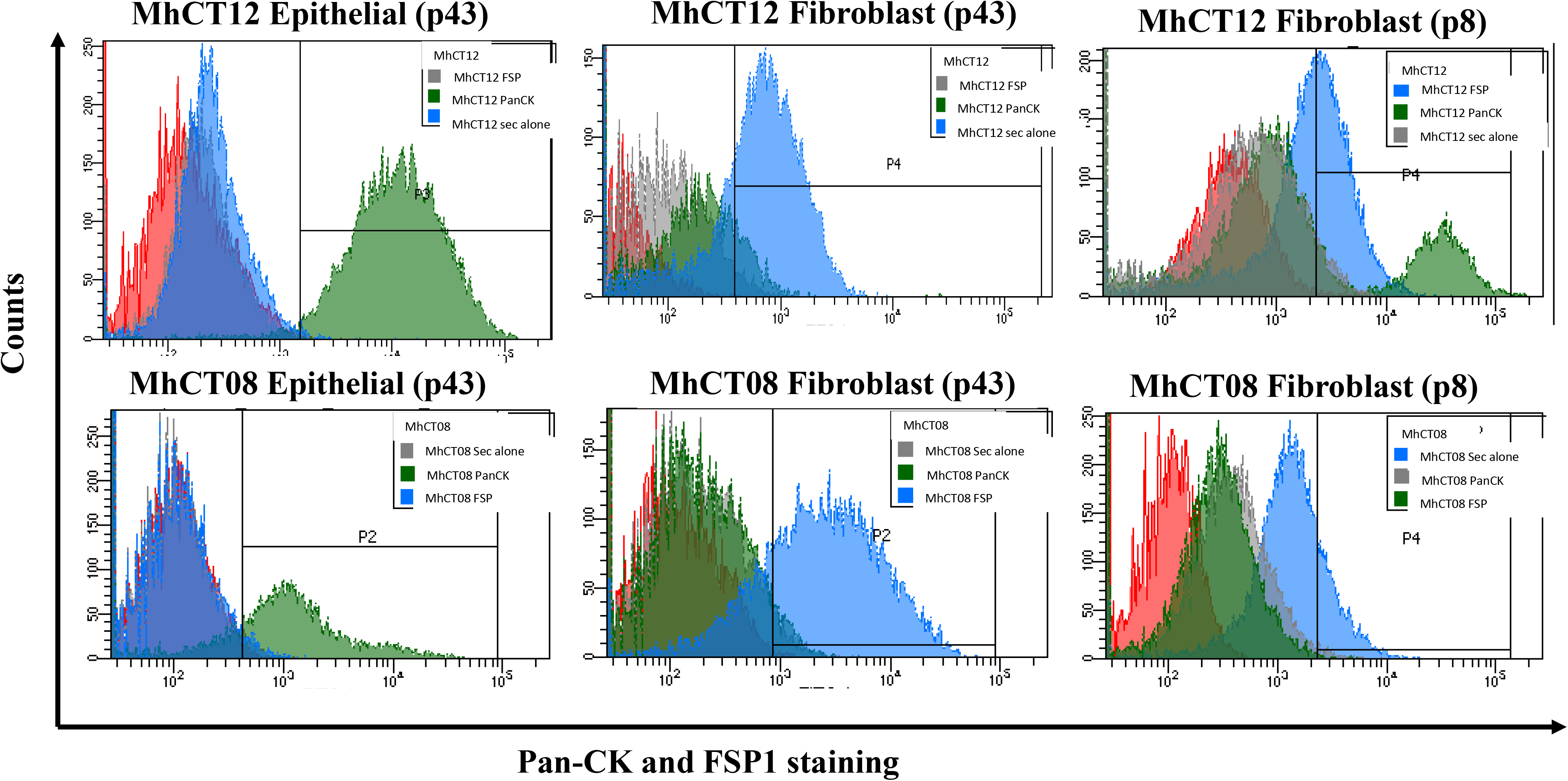
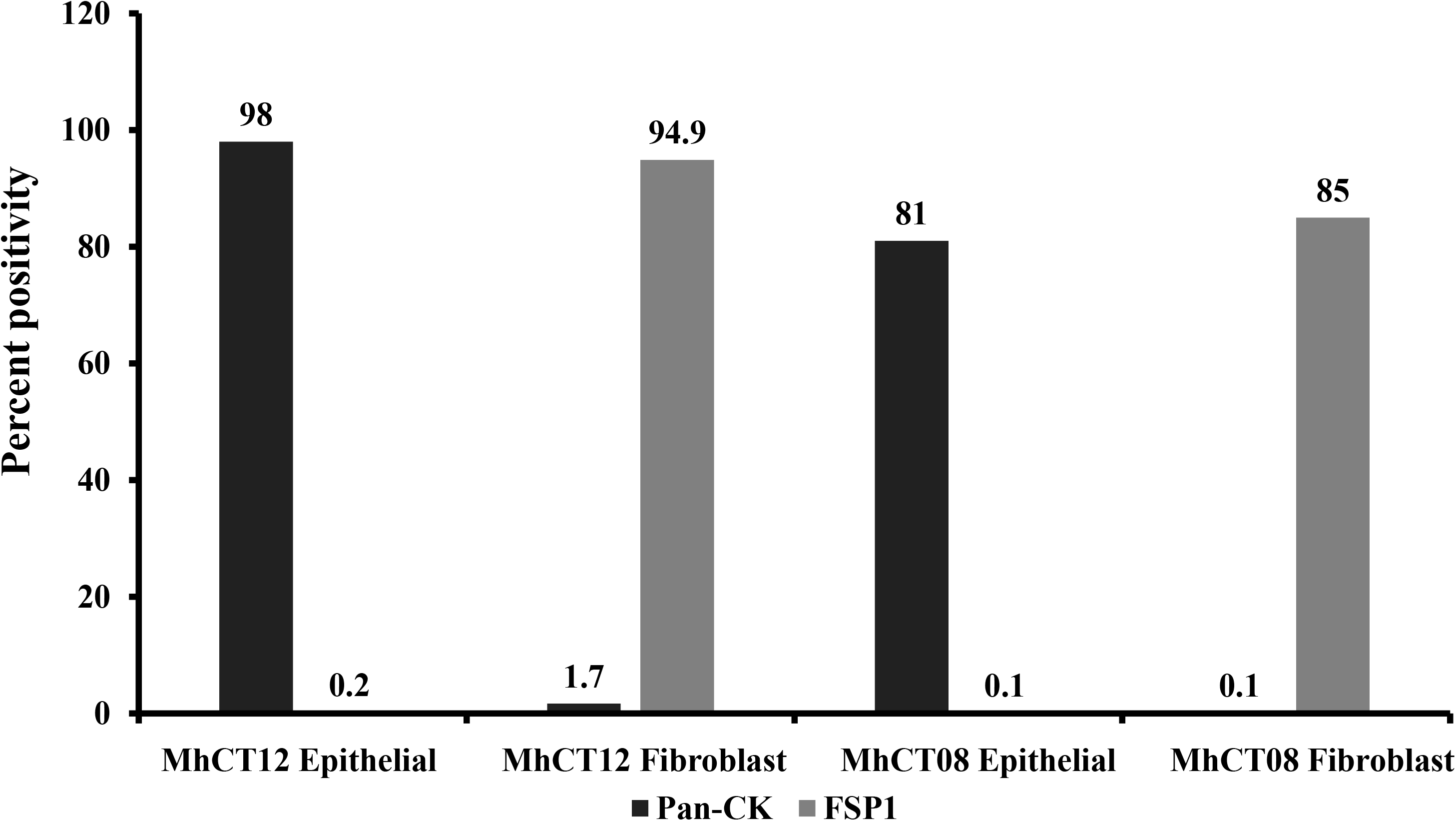
Determination of Purity of Cell lines. (A) Microscopic image analysis. Scale bar - 100μm. (B) Flow cytometry analysis using PanCK and FSP1. (C) Comparison between the primary cultures PanCK and FSP1 positivity.

### Purity

The epithelial and cancer associated fibroblast nature of the cell lines was confirmed by the presence of surface markers, using flow cytometry. The removal of fibroblasts from epithelial population was confirmed by negative staining with FSP-1 antibody (20,25). Similarly, a pure population of fibroblast was confirmed by negative staining with PanCK antibody (26). As depicted in Fig. 1B, 1C, the cell lines were characterized at both an early (p8) and late passage (p43). MhCT12 and MhCT08 epithelial cells showed 98% and 81% PanCK positivity respectively, with less than 0.2% FSP positivity, confirming that epithelial populations were not contaminated with fibroblasts. Similarly, MhCT12 and MhCT08 fibroblasts showed 94.9% and 85% FSP-1-positivity respectively. Both the fibroblast populations did not show any significant PanCK staining thus confirming their purity (Fig. 1C). In addition, the respective epithelial and fibroblast cell lines were stained both at early passage and later passages. As evident from Fig. 1B, primary cultures at later passages showed exclusively PanCK and FSP-1 enriched populations as compared to the early passage.

### HPV detection

As depicted in Fig. S1A, immunocytochemistry analysis revealed that both the epithelial and fibroblast cells were lightly stained for p16 and E7. P16 antibody stained the nuclear region thus confirming the HPV positivity of the established lines (Fig. S1A). Fragments of 450bp were amplified by using MY09/MY11 primers thus confirming the HPV positive status of the established primary cultures (Fig. S1B).

### Ploidy determination

The DNA content of the primary cultures was determined by performing ploidy analysis. Normal human lymphocytes were taken as a control for diploid DNA content. The mean channel of cells that were in the G0 phase was divided by that of the lymphocytes to determine DNA index of each cell line. The DNA indices of MhCT08 and MhCT12 cells were 1.1 and 0.8 respectively (Fig. 2A). The results therefore indicate that both the patient samples have abnormal DNA content which might be responsible for the immortalization of these cells.

**Figure 2.**
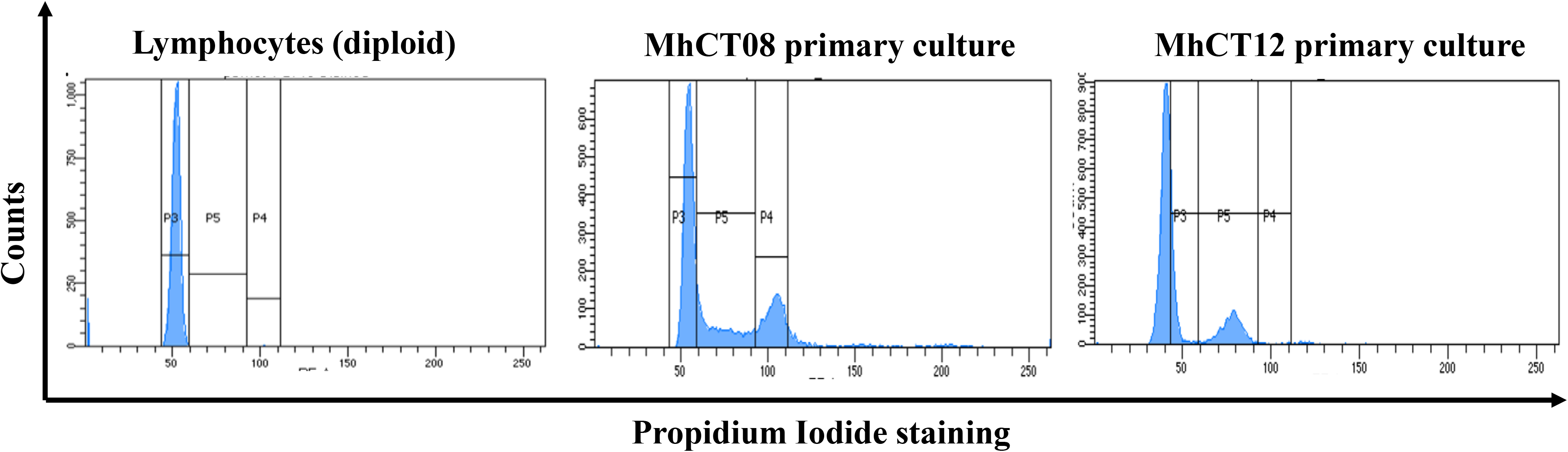
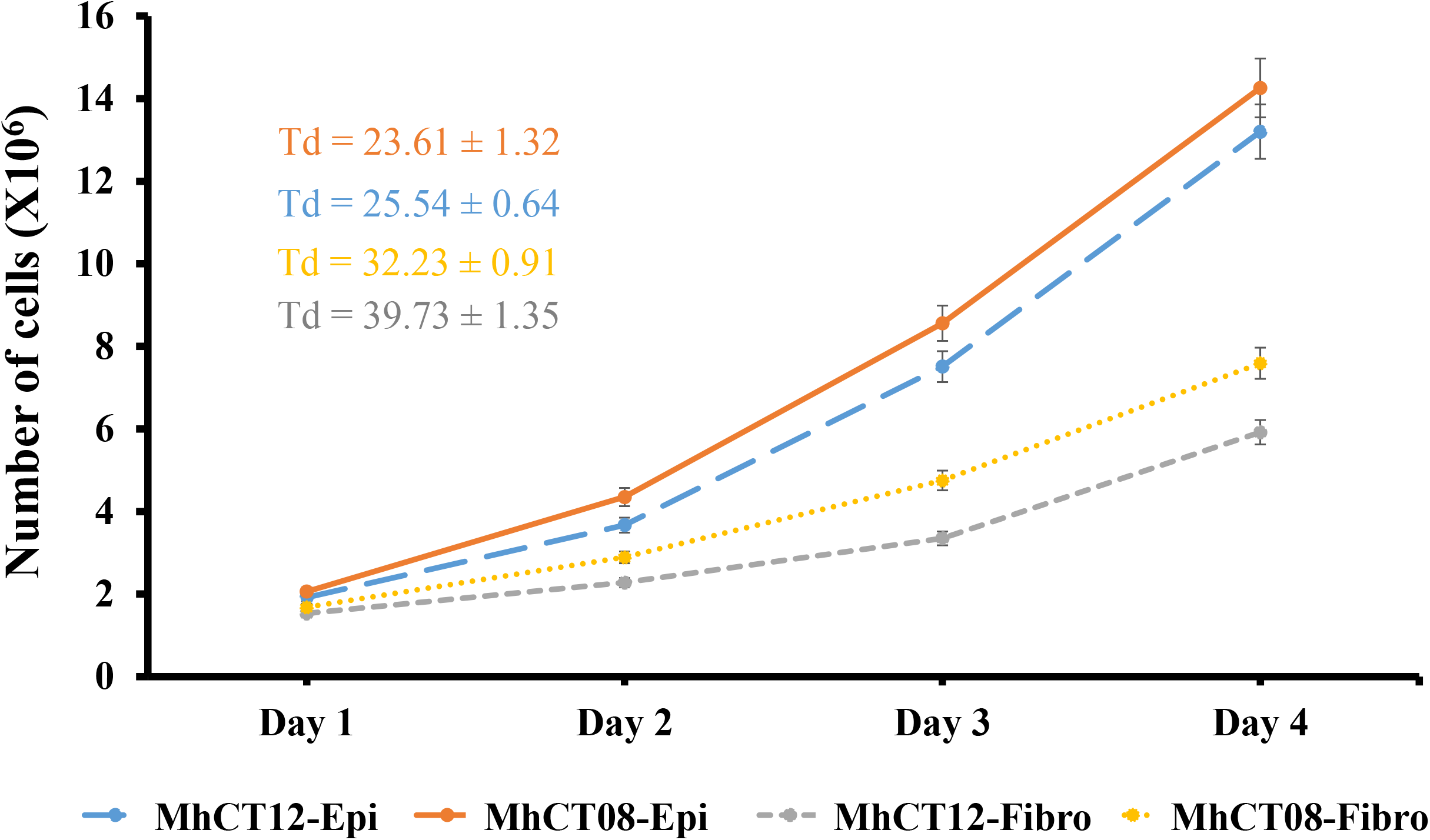
Characterization of Established Cell Lines. (A) Ploidy determination. Flow cytometric analysis to measure the genomic DNA content of MhCT08 and MhCT12 primary cultures. Normal human lymphocytes were used as a control for diploid DNA content (B) Doubling time of established primary cultures. Td – doubling time

### Growth characteristics

The cell lines exhibited different doubling times as depicted in the Fig. 2B. Doubling time of 23.61 hours and 25.54 hours was observed for MhCT08 epithelial and MhCT12 epithelial cell lines respectively. Both the fibroblast cell lines had higher doubling times than epithelial cells as depicted in Fig. 2B with MhCT12 fibroblast doubling at every 39.73 hours and MhCT08 fibroblast doubling every 32.23 hours.

### STR profiling

The STR profile of the established cell lines was distinct from that of any other cell lines deposited in ATCC and Expasy Cellosaurus STR (CLASTR) database (Table II). These results thus indicated that both MhCT08 and MhCT12 fibroblast and epithelial cells are novel. Similar genotypic identity between epithelial and fibroblast cells from the same patient distinctly proved the autologous nature of the cells (Table III). No cross contamination, especially with Cal27 and HSC3 cell lines (simultaneously handled by author ND) was observed as shown in the Table III.

**Table II.**
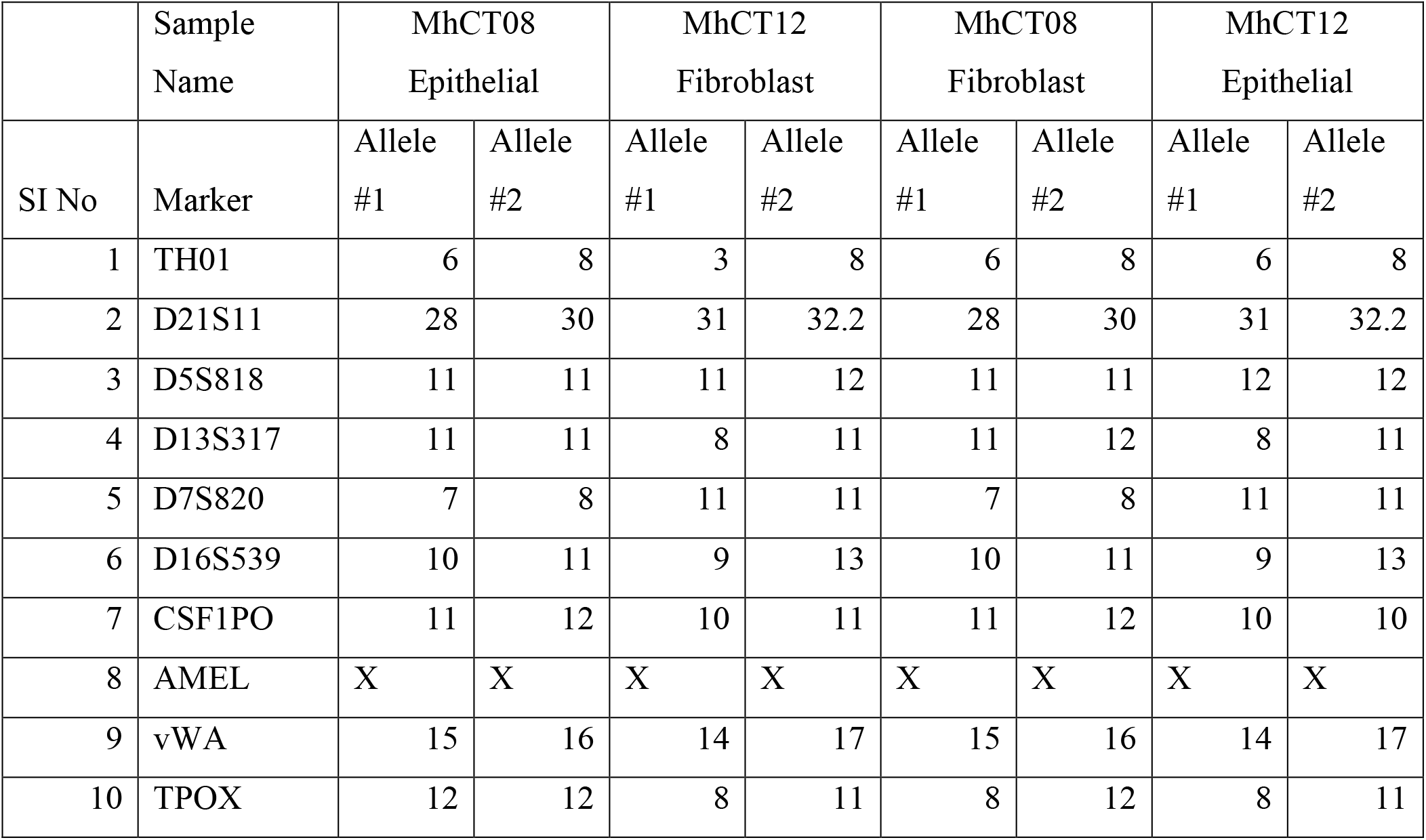
STR profiles of the established cell lines

**Table III.**
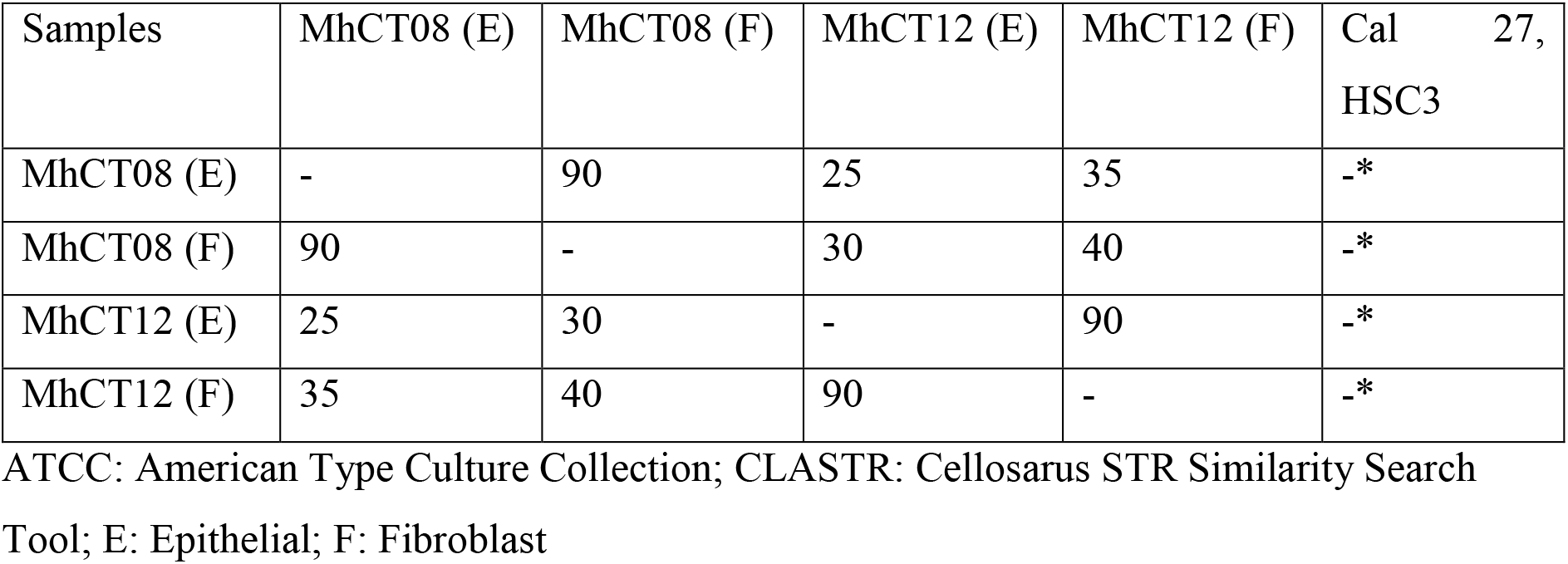
Proof of novelty by STR (* <50% match cut-off from ATCC and CLASTR public databases)

#### Tumorigenic properties of primary epithelial cells

Furthermore, to assess the tumorigenic potential of the epithelial cell lines, various assays were performed post treatment with CAF conditioned medium.

### Proliferation

CAF conditioned medium was taken neat and supplemented with twenty and fifty percent of fresh complete RPMI medium, in which the epithelial cells were allowed to grow till 72 hours. Proliferative potential of the cells was calculated every 24 hours. Epithelial cells showed a significant increase in proliferation when treated with CAF conditioned medium. Since no significant difference was observed between neat and supplemented conditioned mediums, only neat conditioned medium was taken for subsequent experiments. A slight decrease in proliferation was observed at 72 hours, which may be attributed to over-confluency of the cells leading to floating cells. Maximum proliferation under the effect of neat CAF conditioned medium was observed at 48 hours for both the cell lines (Fig. 3A and 3B).

**Figure 3.**
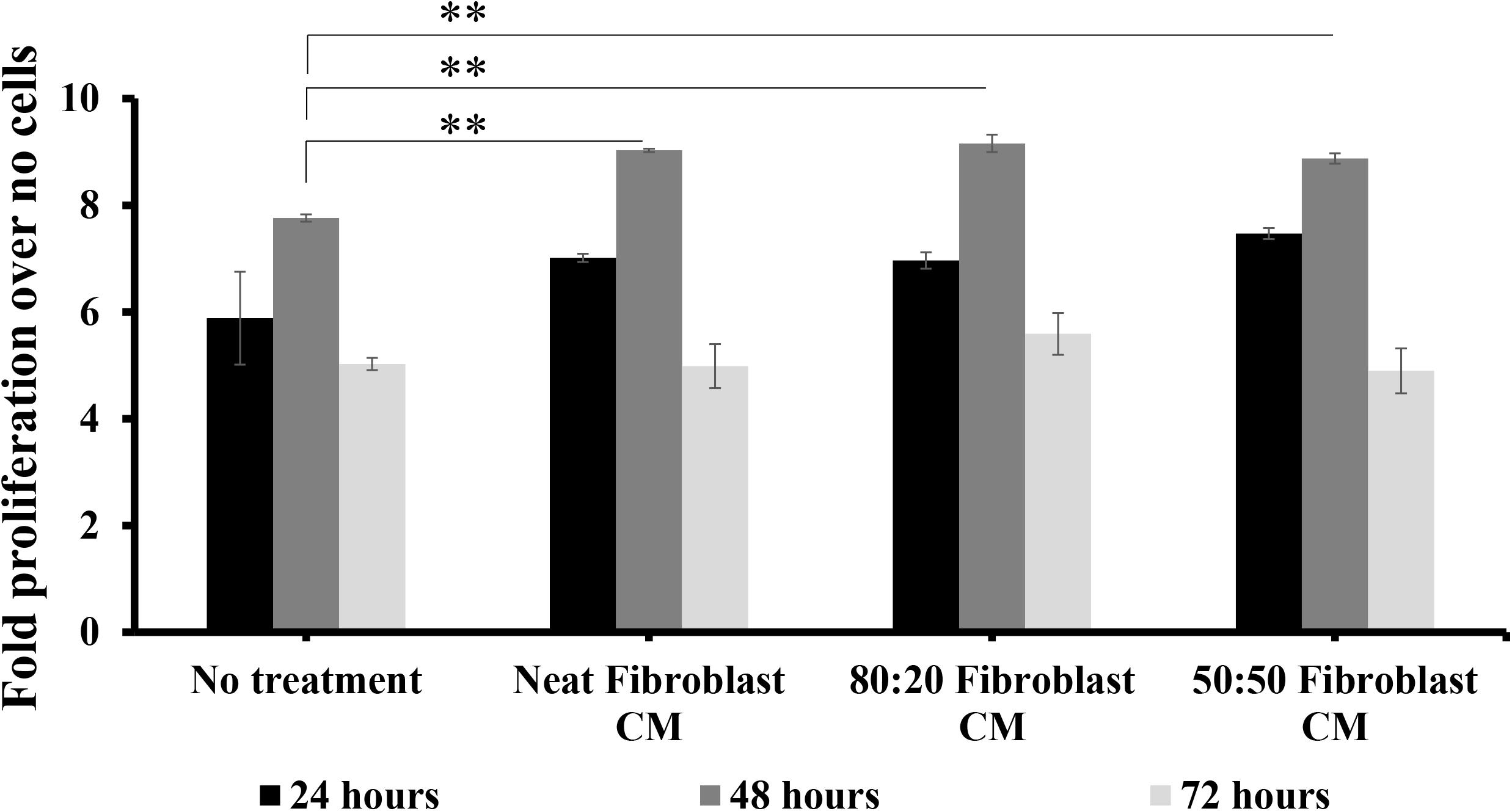
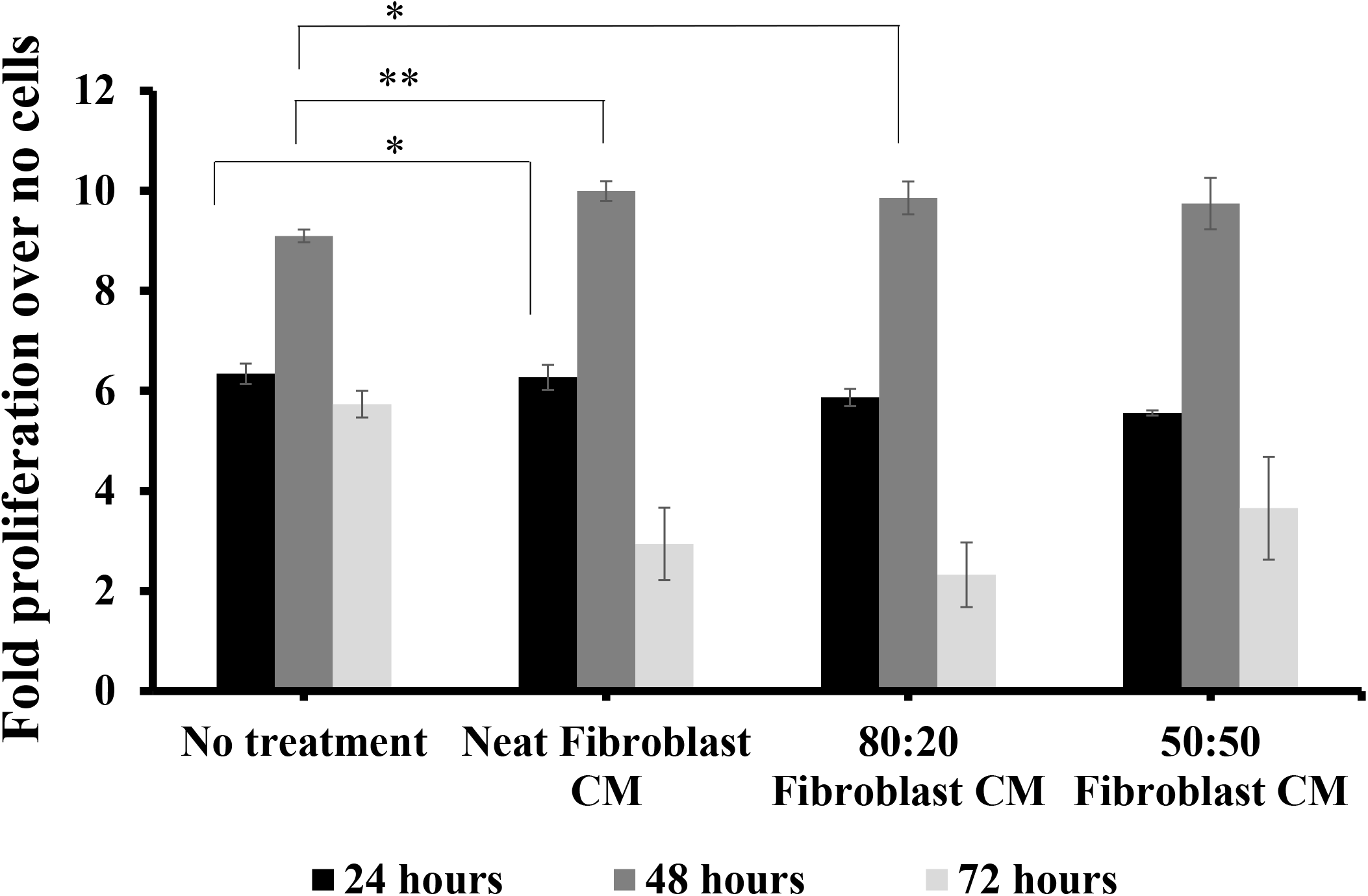
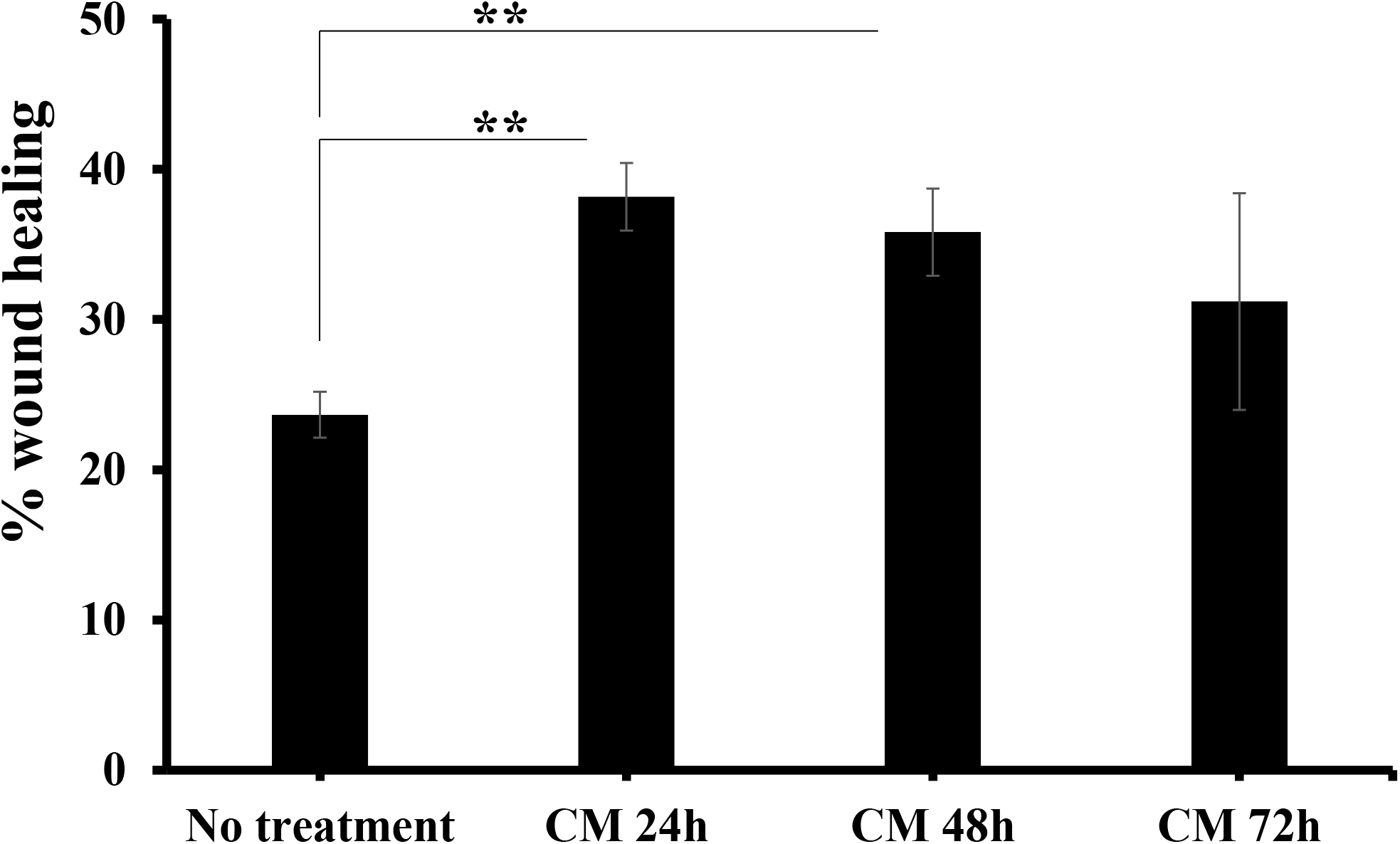
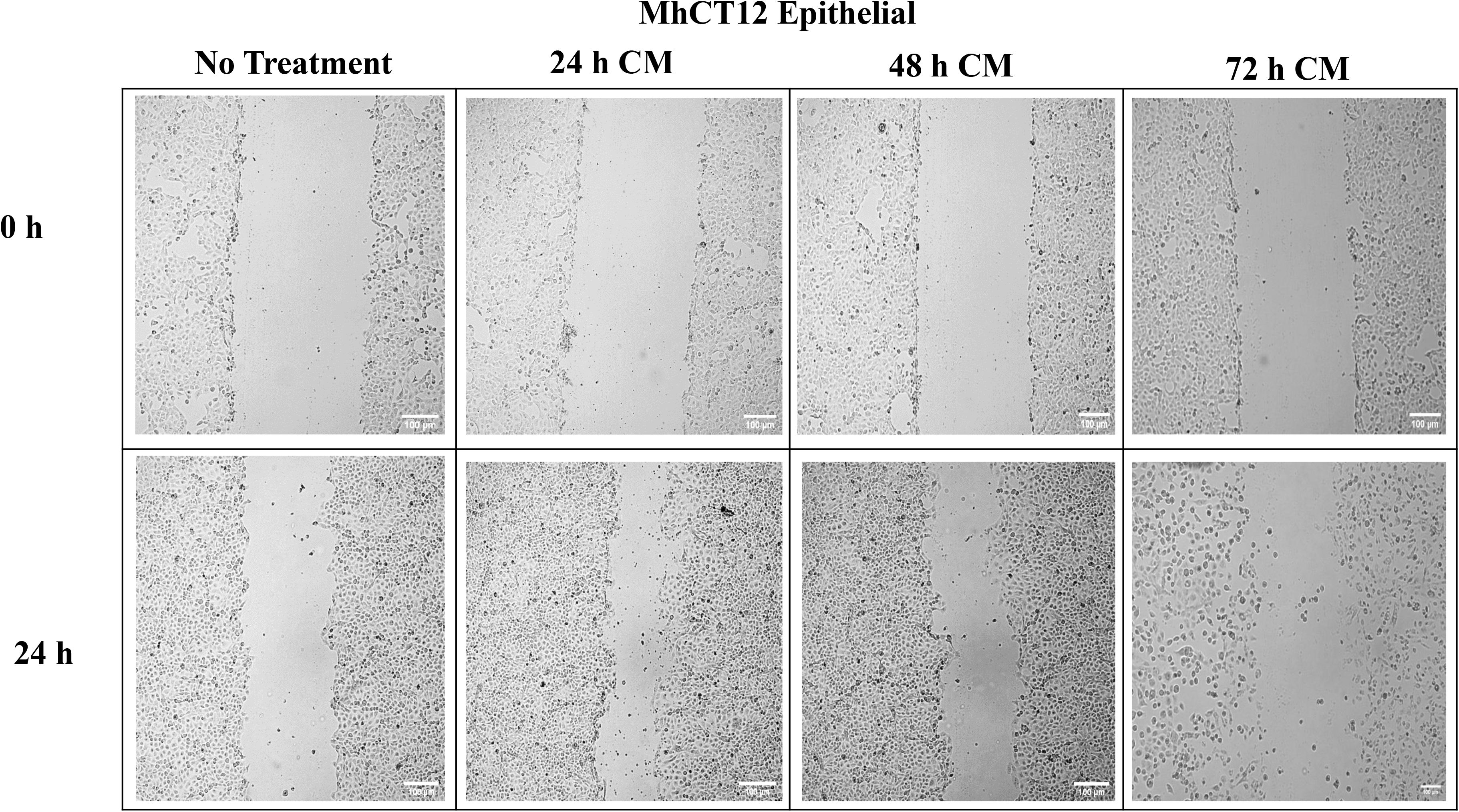
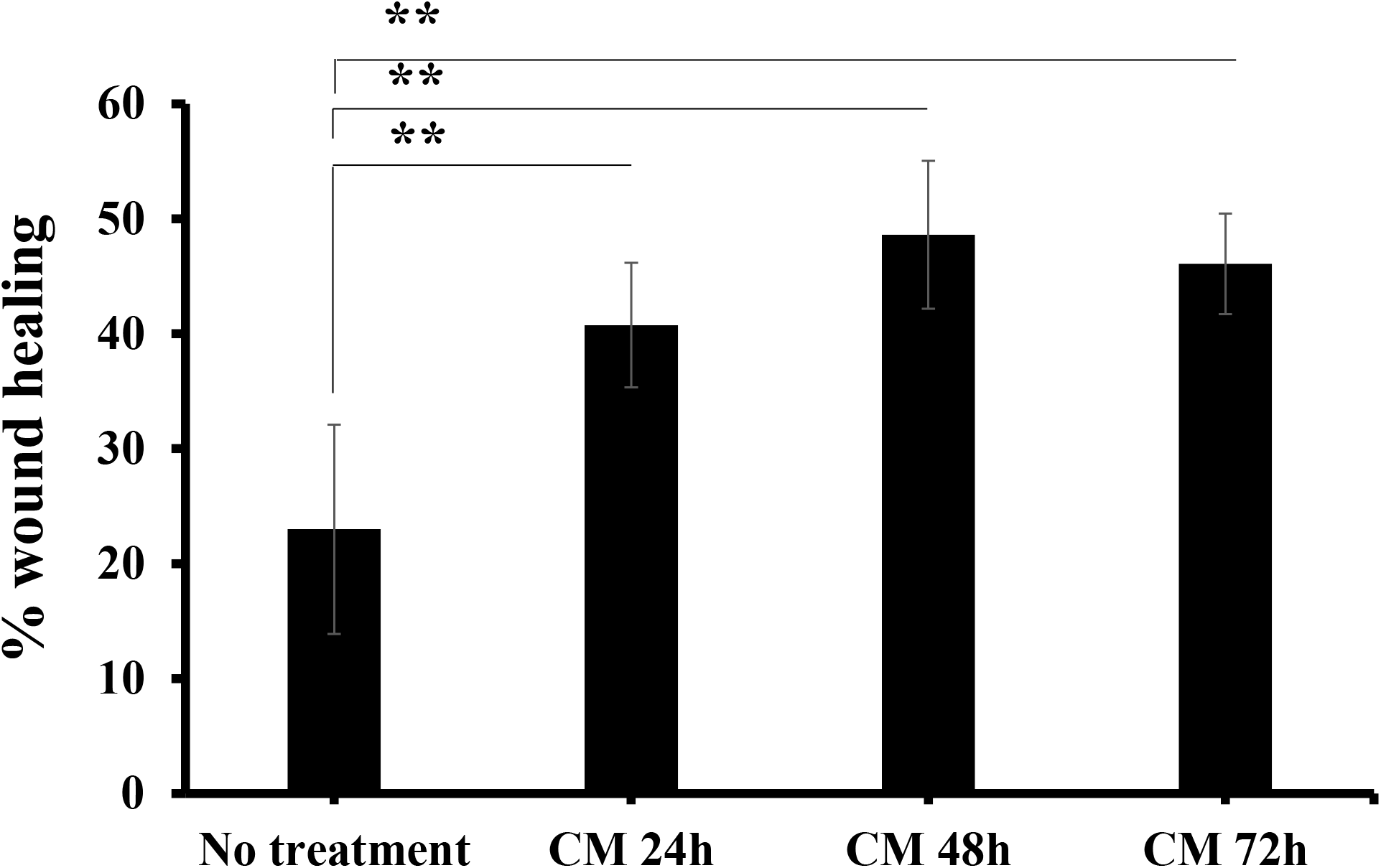
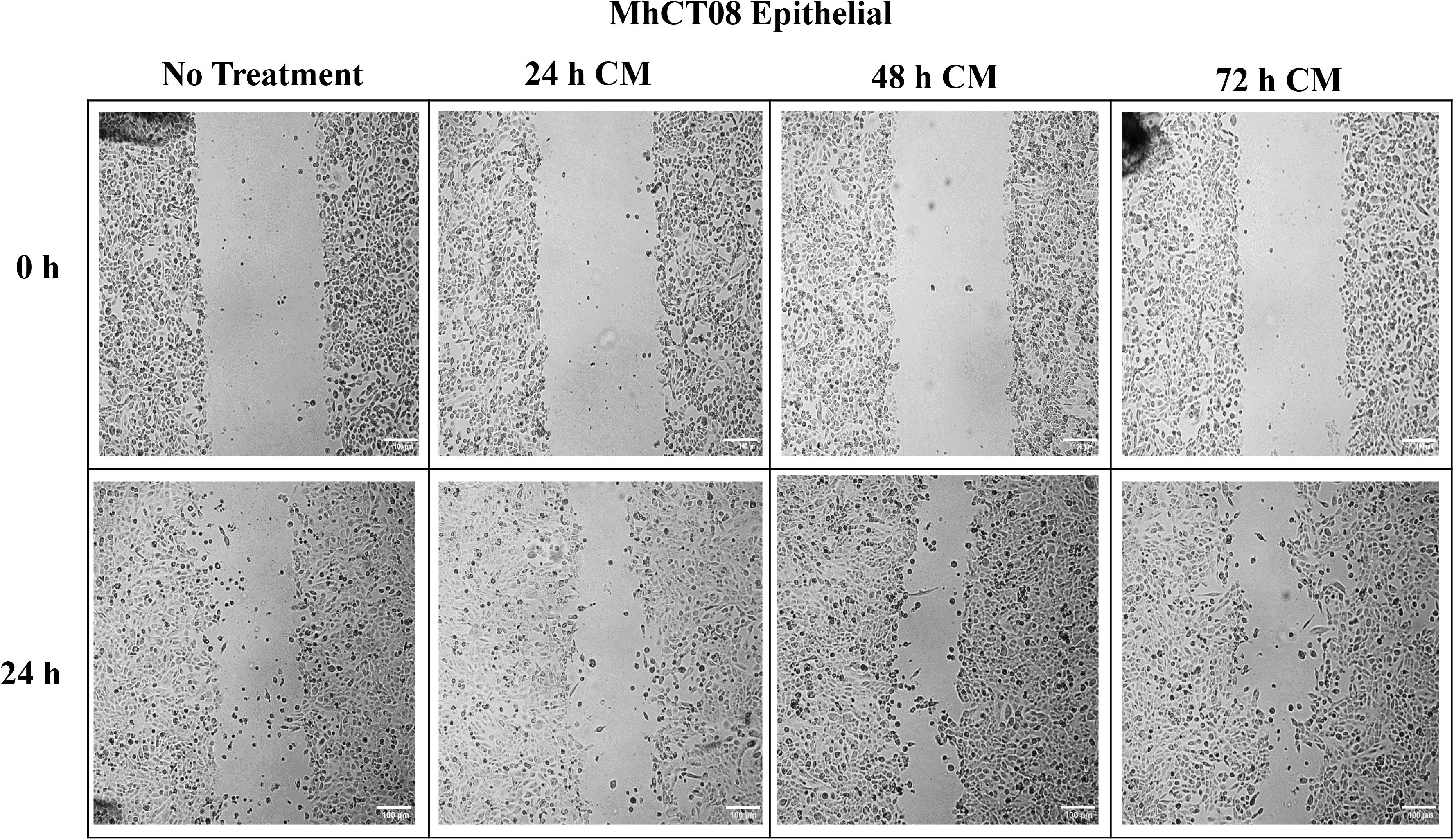
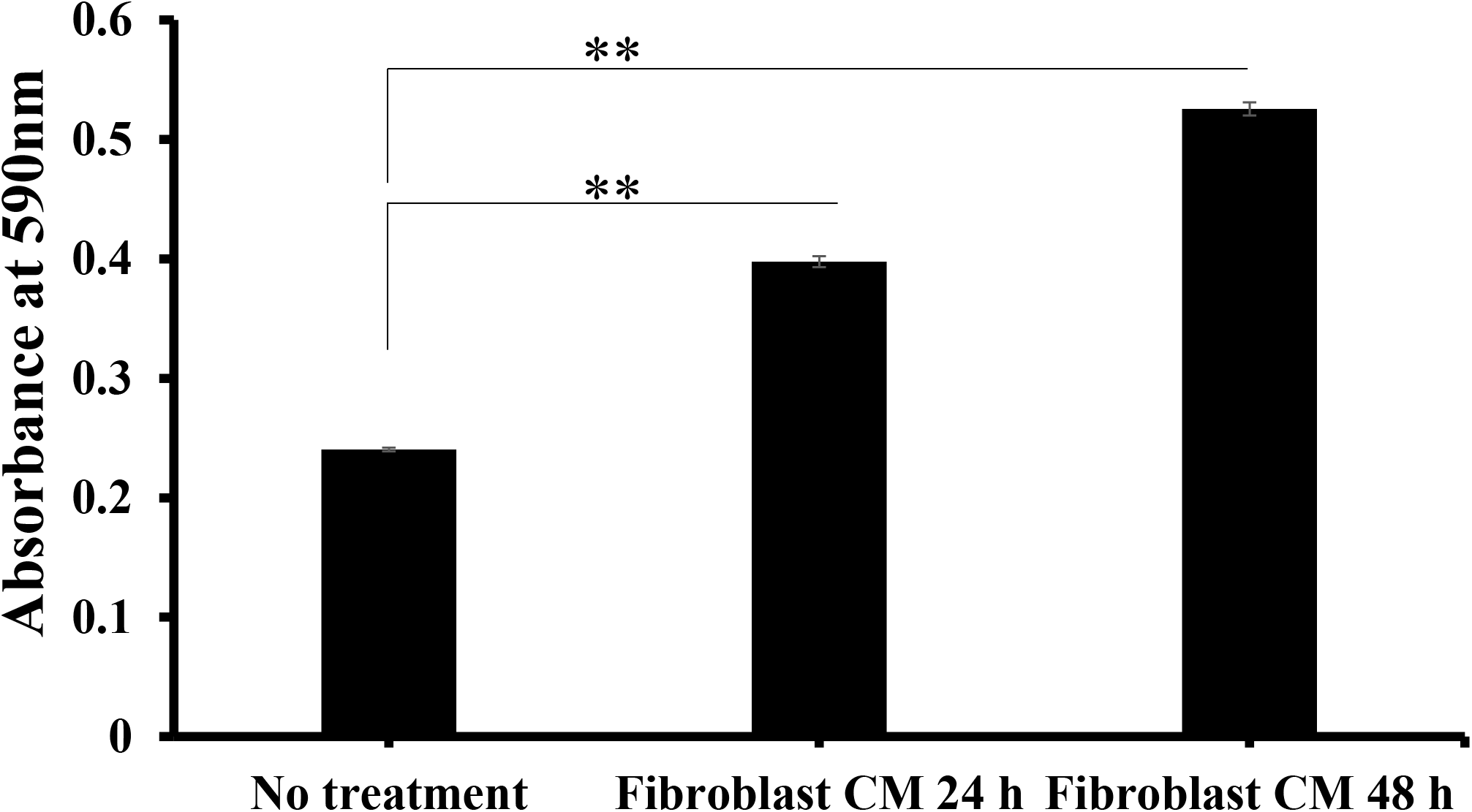
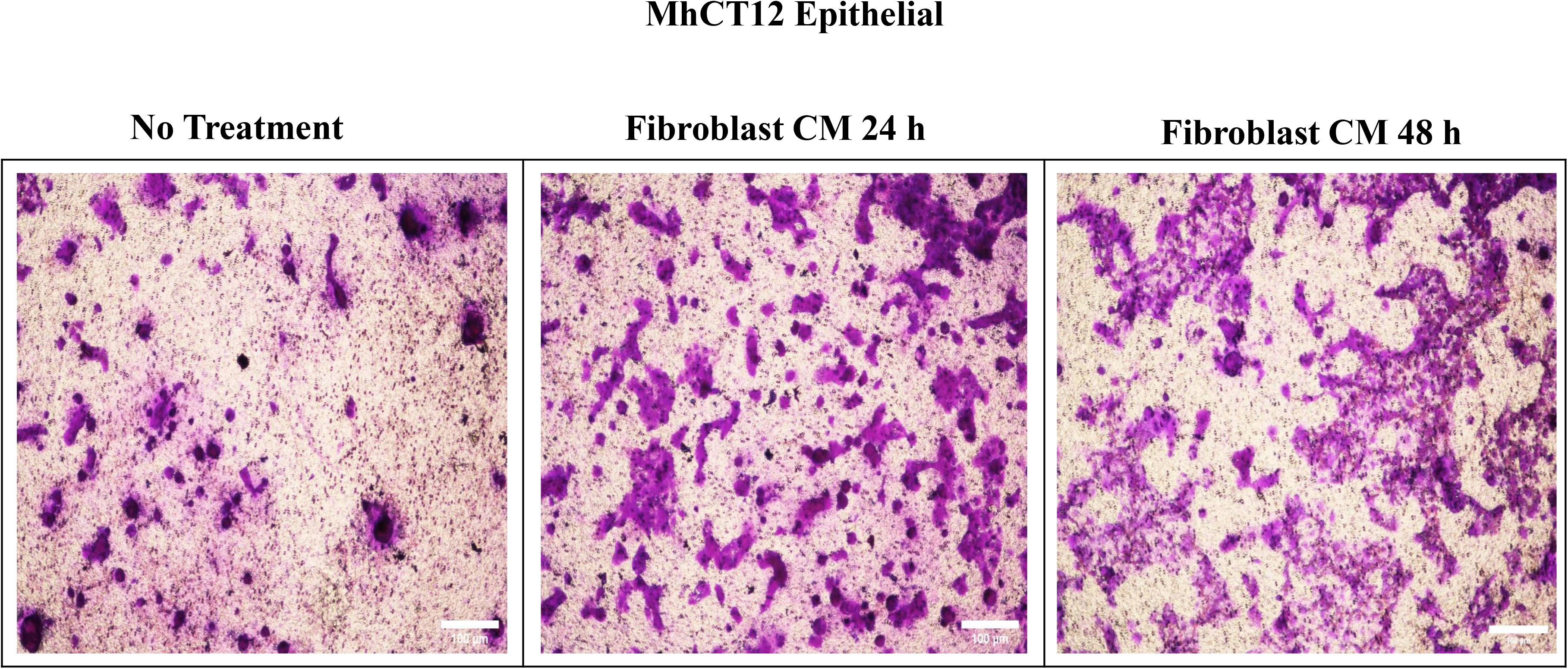
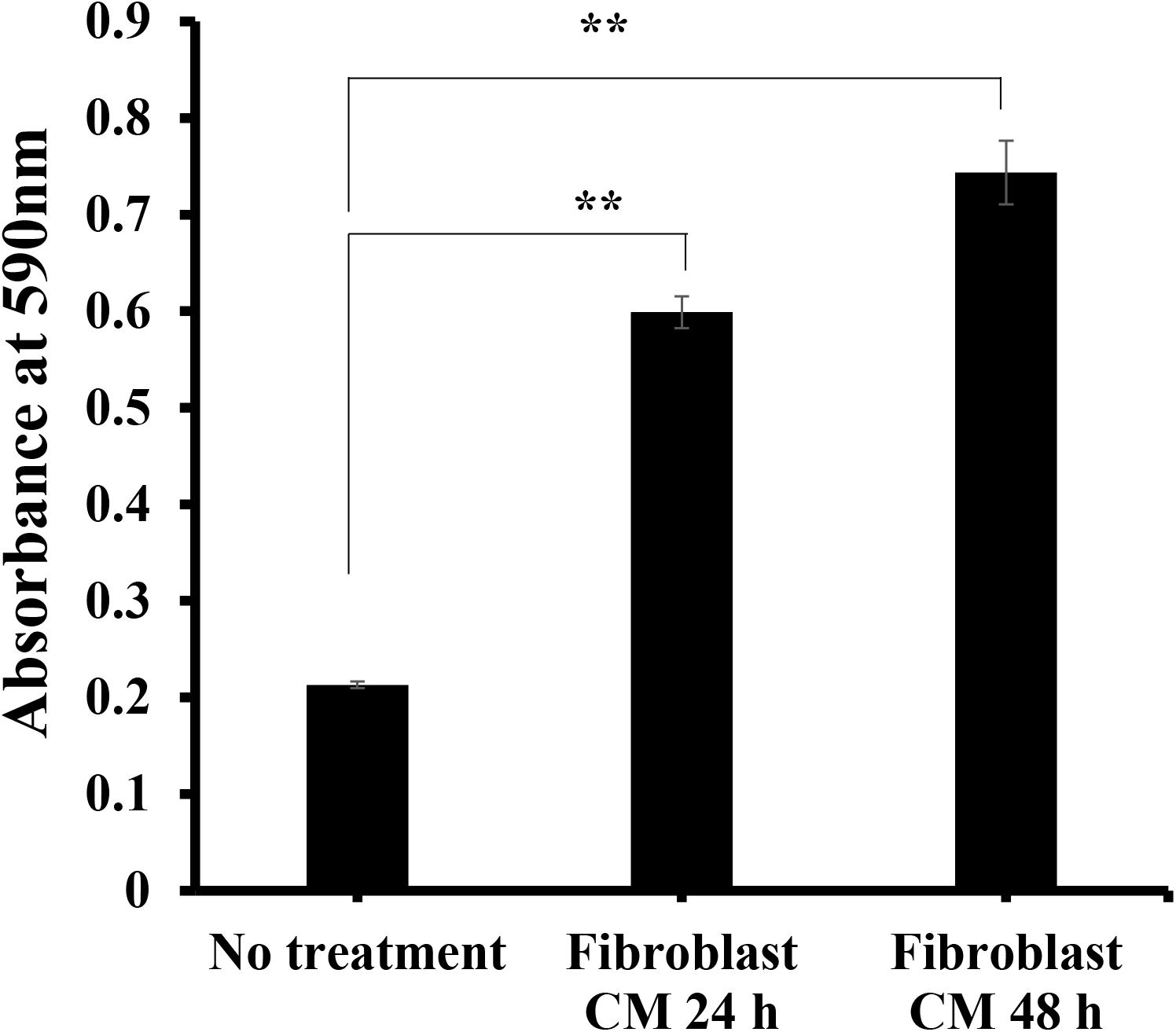
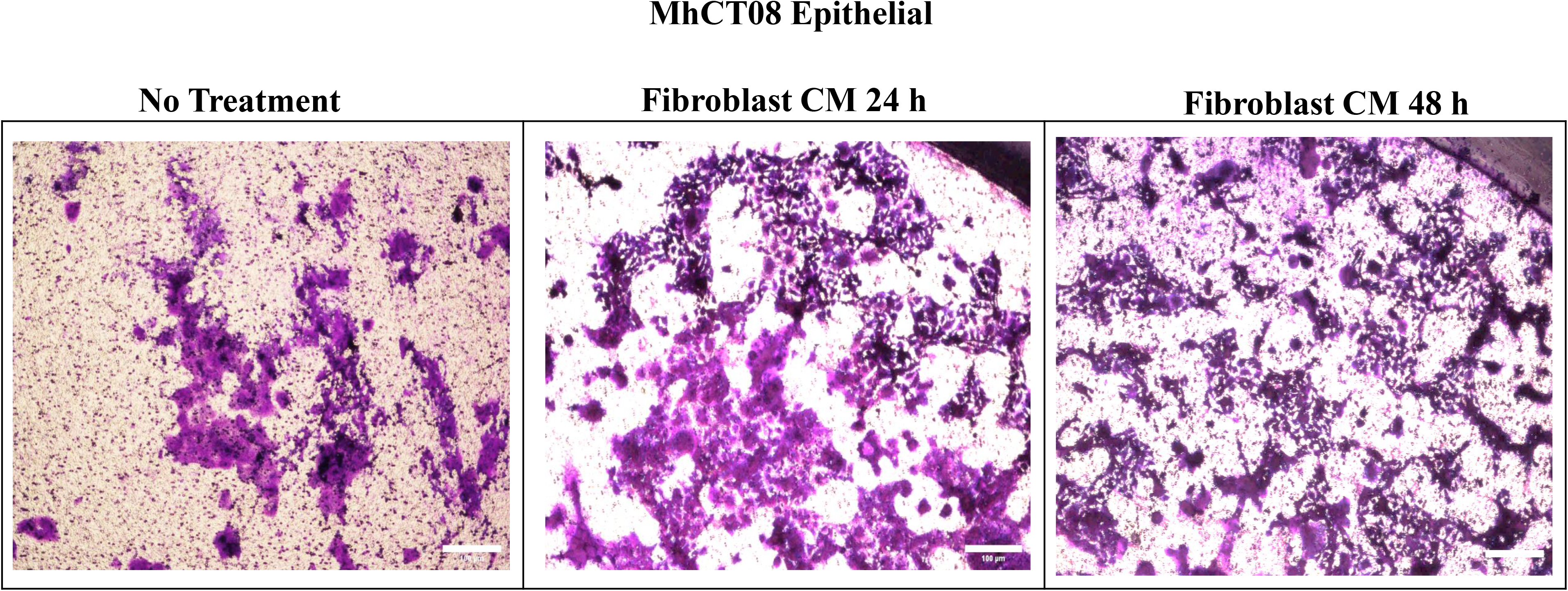
Tumorigenic properties of Established Cell Lines. (A-B)Proliferation of indicated epithelial cells under the effect of fibroblast conditioned medium at different time points. (C-D) Percentage wound closed represented graphically and pictorially for MhCT12 epithelial cell line under the effect of fibroblast conditioned medium collected at indicated time points (scale bar 100μm). (E-F) Percentage wound closed represented graphically and pictorially for MhCT08 epithelial cell line under the effect of fibroblast conditioned medium collected at indicated time points (scale bar 100μm). (G-H) Invasive potential as measured by crystal violet staining represented graphically and pictorially for MhCT12 epithelial cell line under the effect of fibroblast conditioned medium collected at indicated time points (Scale bar 100μm). (I-J) Invasive potential as measured by crystal violet staining represented graphically and pictorially for MhCT08 epithelial cell line under the effect of fibroblast conditioned medium collected at indicated time points (Scale bar 100μm). Abbreviations: CM – conditioned medium. Neat – Complete conditioned medium; 80:20 – 80% conditioned medium + 20% complete fresh medium; 50:50 – 50% conditioned medium + 50% complete fresh medium. (**p<0.01, * p<0.05)

### Wound healing assay

Treatment with neat conditioned media from the autologous fibroblast pair significantly increased the migration potential of MhCT12 epithelial from 23% to 38% (Fig. 3C) and 22% to 48% for MhCT08 epithelial (Fig. 3E) (p-value < 0.05) as depicted in Fig. 3D and 3F. Treatment with CAF conditioned medium significantly increased the wound healing potential of their respective epithelial counterparts

### Invasion assay

Treatment with conditioned media from the autologous fibroblast pair significantly increased the invasive potential of both MhCT08 and MhCT12 epithelial cell types (p-value < 0.05) as depicted in Fig. 3G – 3J.

### Sphere formation assay

Treatment with conditioned media from the autologous fibroblast pair significantly increased the sphere formation potential of both MhCT08 (n = 60, p<0.01) and MhCT12 epithelial (n = 35, p<0.05) cell types, as compared to no treatment conditions of MhCT08 (n = 36) and MhCT12 (n = 24) as depicted in Fig. 4. Treatment with MhCT08 fibroblast conditioned medium however influenced the epithelial cells to form bigger spheres (5.4 × 10^7^, p<0.01) as compared to the treatment with MhCT12 fibroblast conditioned medium. (1.2 × 10^7^, p<0.01)

**Figure 4.**
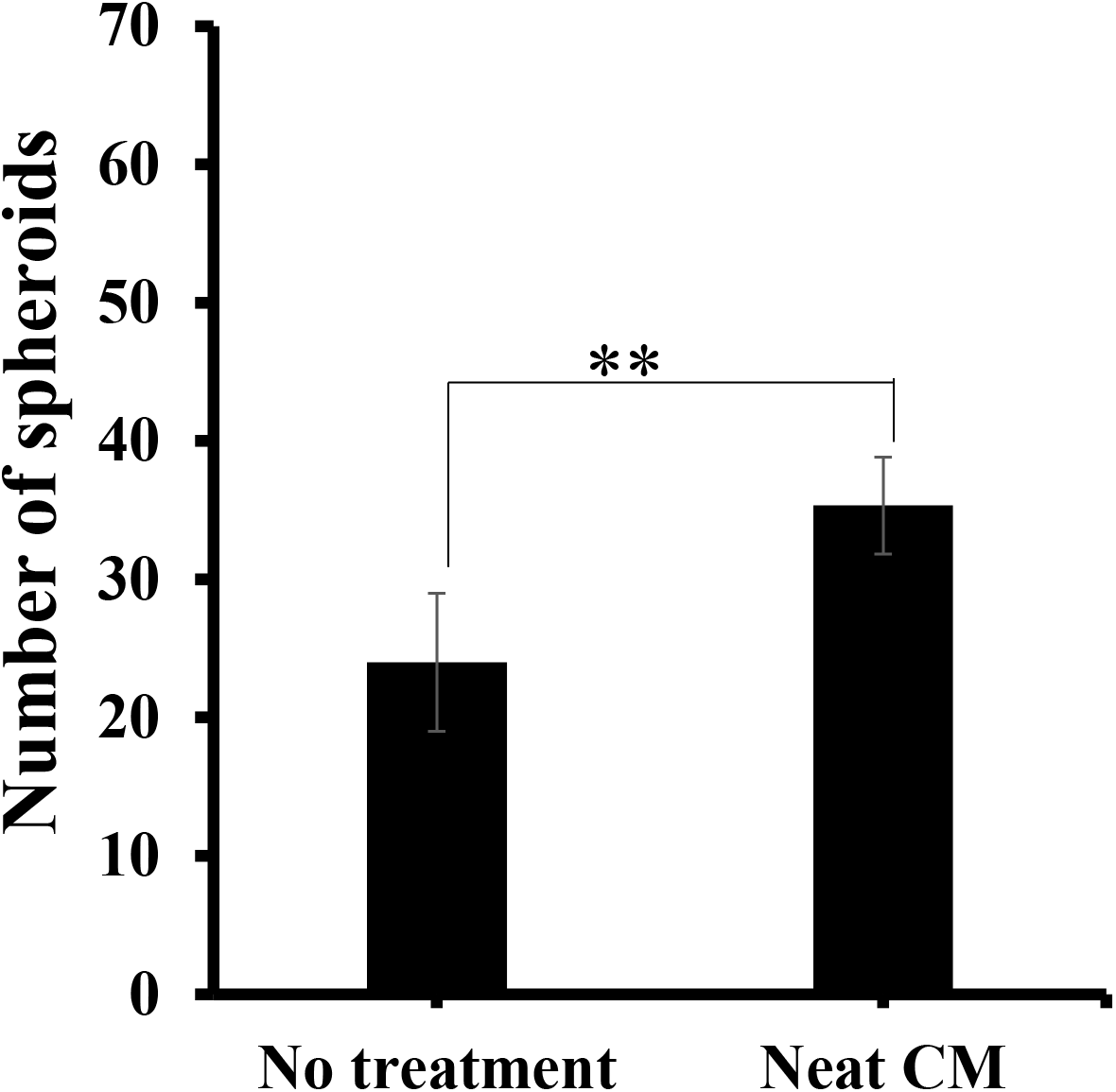
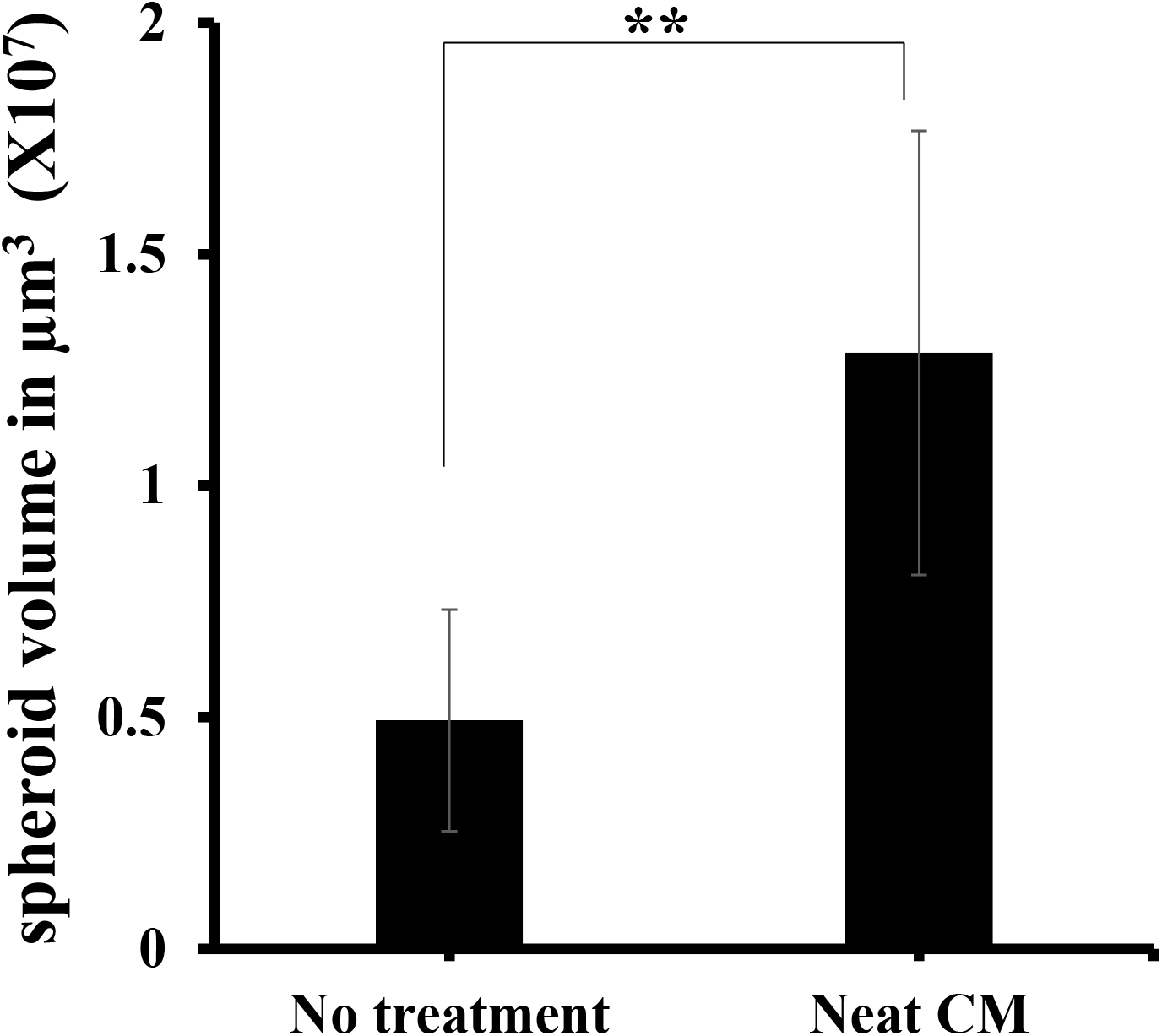
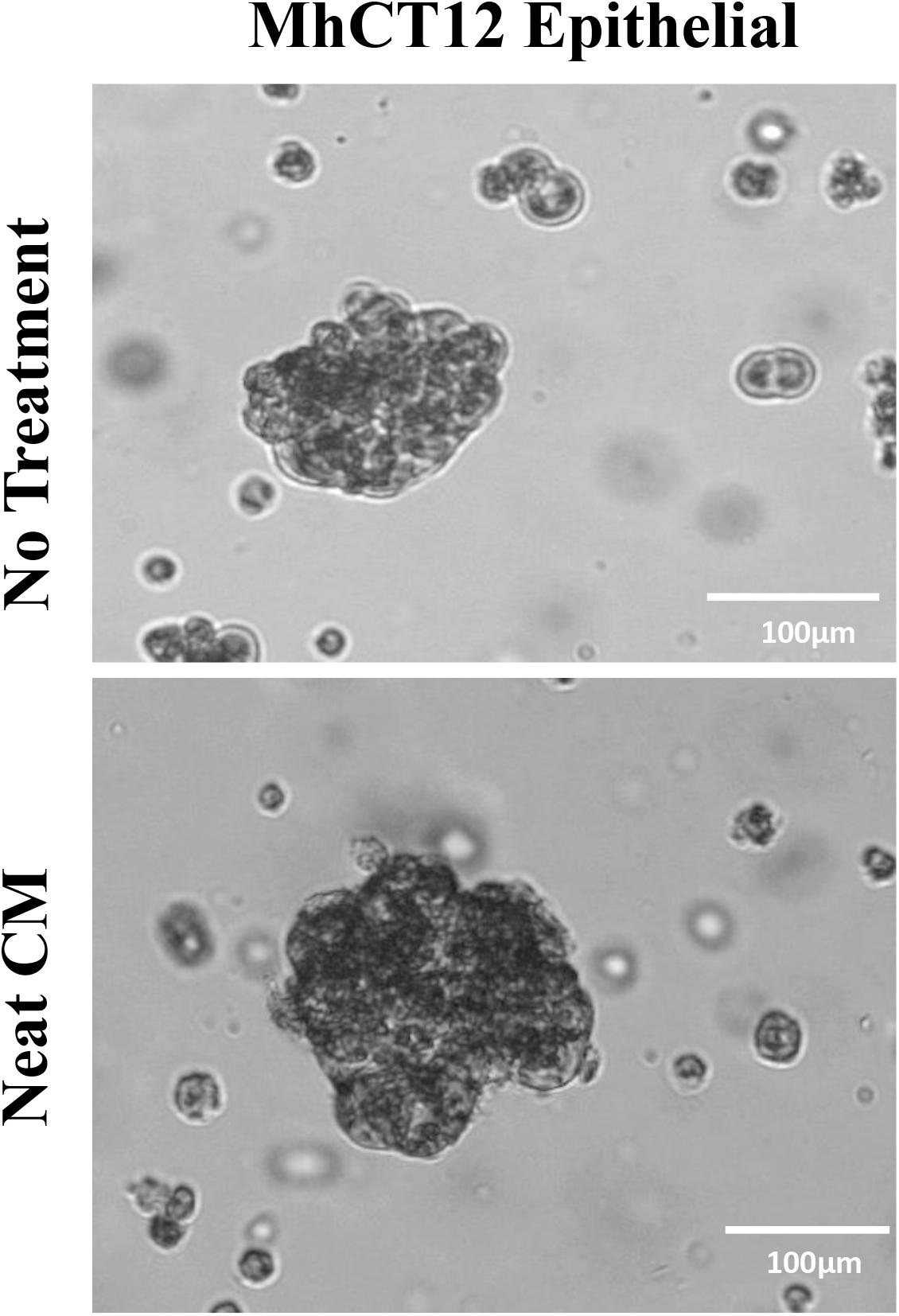
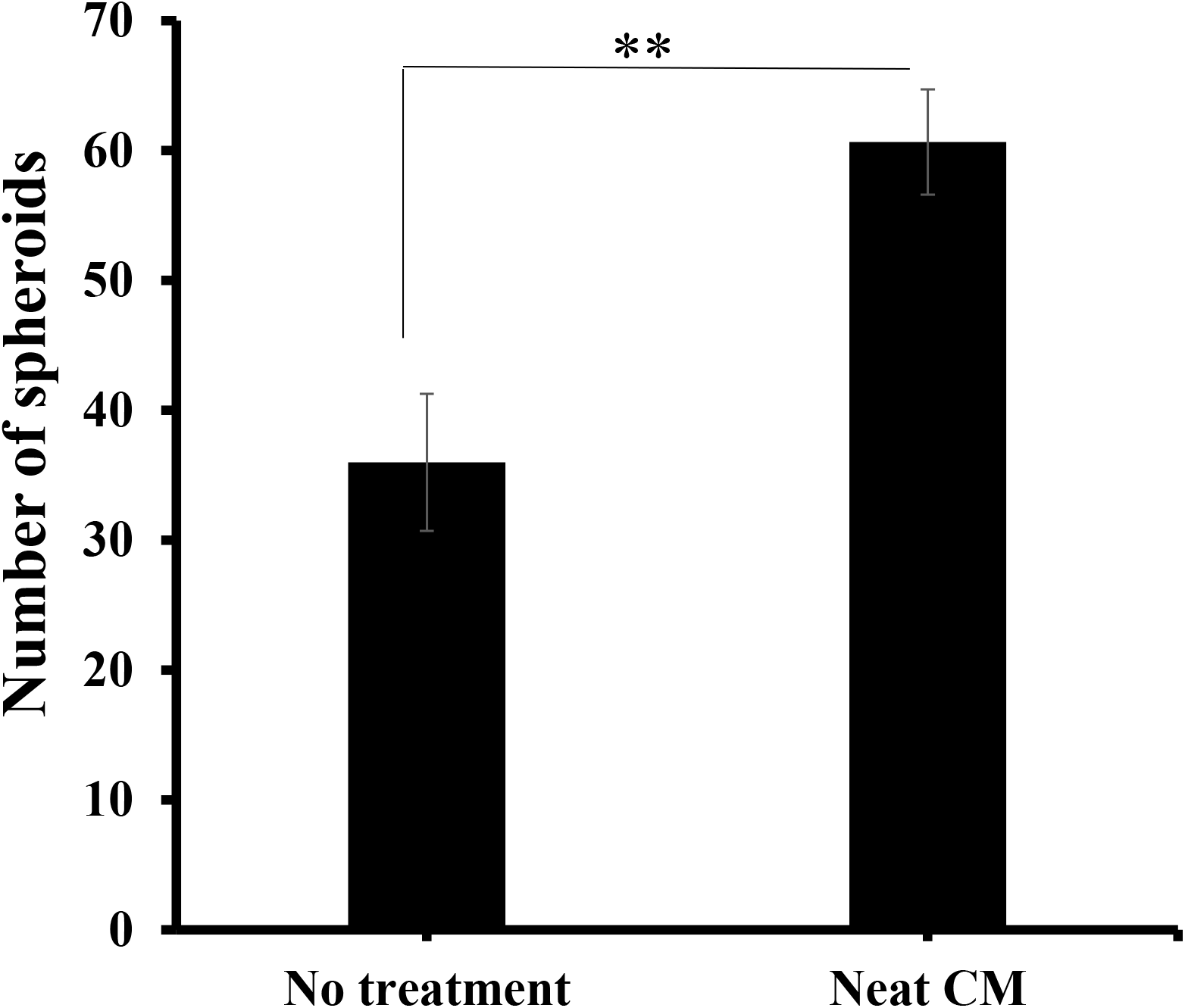
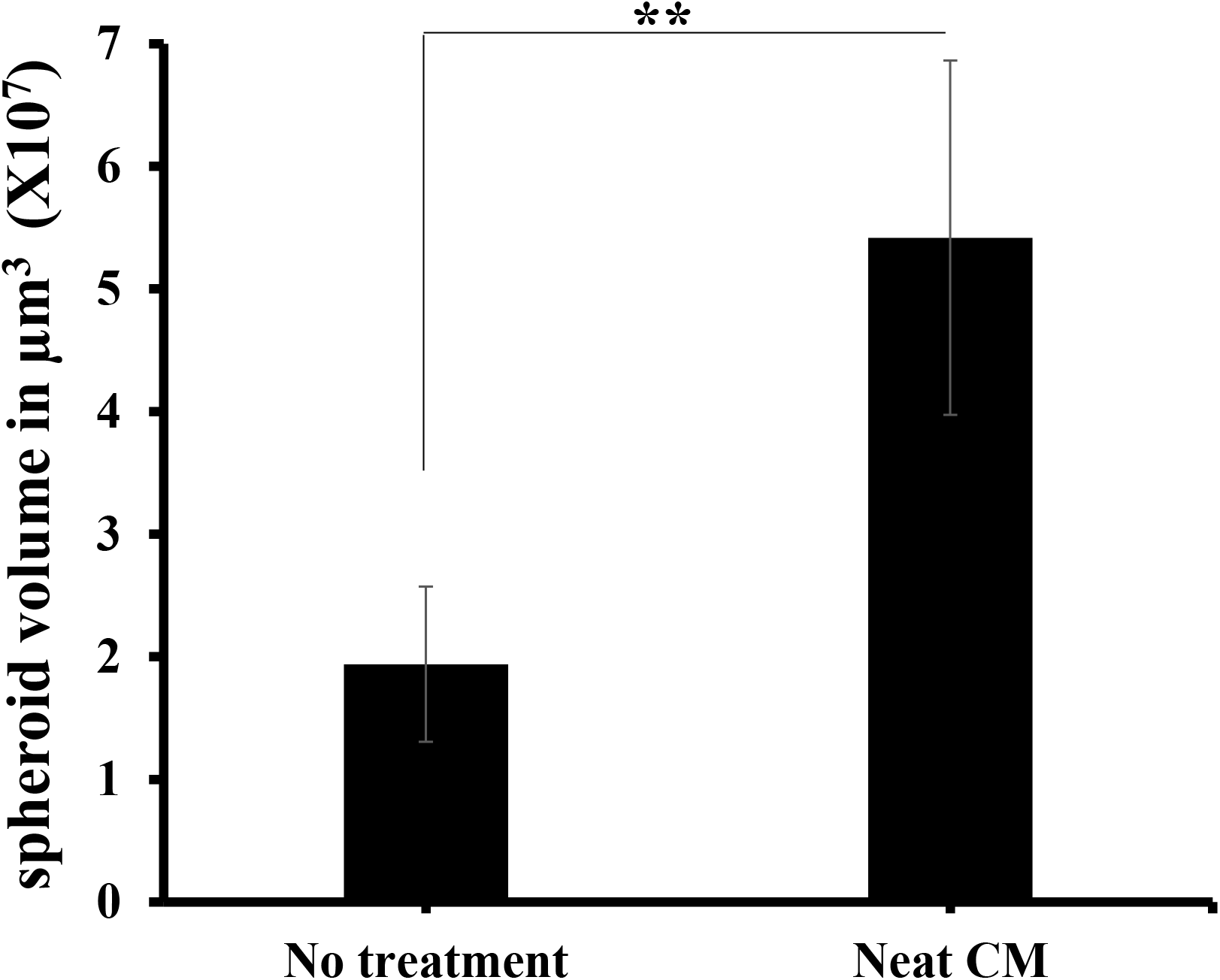
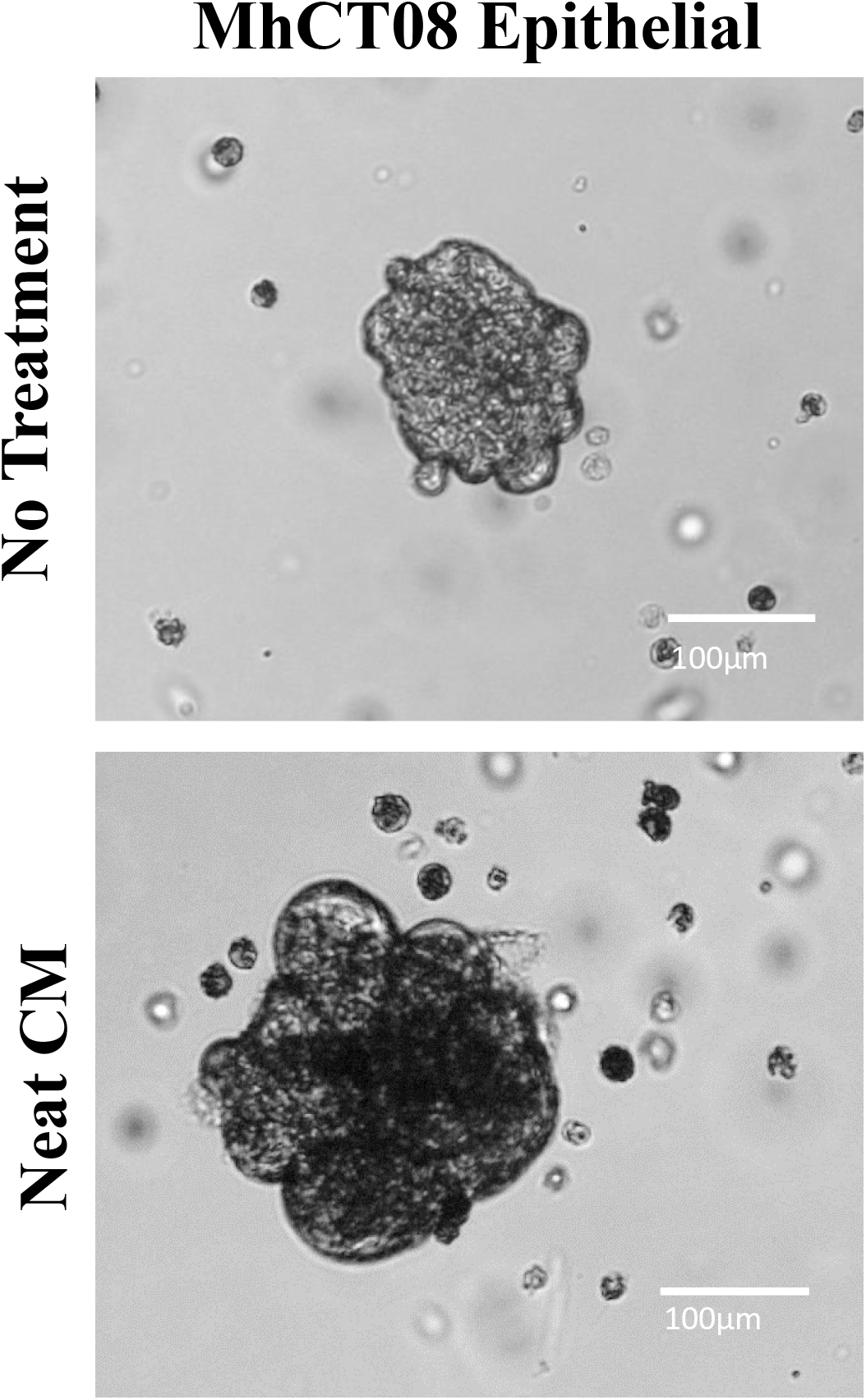
Sphere formation assay. (A-C) Sphere formation potential of MhCT12 and (D-F) MhCT08 under the effect of fibroblast conditioned medium (scale bar 100μm). CM– Conditioned medium. (**p<0.01, * p<0.05)

## Discussion

The present study details the establishment and characterization of two novel autologous epithelial and CAF cultures from two 65 years old Indian females without any risk habits, diagnosed with squamous cell carcinoma of the tongue.

Like in any other cancer, in vitro cell based models from oral cancer have proven to be an essential tool to enhance the current understanding of tumor progression, tumor heterogeneity and stroma–tumor cross talk. Establishment of various cell lines excised from patient tumor have been a long-known practice to study the etiology of the tumor. A large number of studies have also reported the importance of tumor stroma in cancer progression (27–29). Various articles report the establishment of epithelial cell lines from patient samples. These epithelial cell lines were derived from different tissue origins, for example, buccal mucosa (15), gingivobuccal mucosa (14), oral cavity (30–32), tongue (13,33), sinonasal (34), pyriform fossa (35), lower alveolus and retromolar trigone (33). To elucidate the impact of different mutations on tumor behaviour, recently 16 cell lines from HNSCC were established by Hayes et.al (36). Notably, only a few oral cancer cell lines have been established from Asian countries, including Indian patient population (30,32,33,35). All the studies reported establishment of the primary cultures from both females and males, without any inclination towards one sex.

Traditional risk factors for the development of oral cancer include chewing, smoking and alcohol consumption. The primary cultures reported were from different stages of tumor and with and without habits. In the present study, both the patients were without any reported risk habits. Overall in India, OSCC cases without any known risk habits are comparatively very less (37–39). Therefore comparing these primary cells with the other established primary cultures, from patients with risk habits, might give insights about etiology of the disease progression in both cases.

Mulherkar et.al described the establishment of NT-8e, an oral squamous cell carcinoma cell line using a mouse xenograft model (35). Certain studies have also reported directly digesting the tumor tissue and seeding the cell suspension (30,33). Majority of reports have described establishing primary cultures by explant culture method (13–15,32,34). The current study also used explant culture method for MhCT12 and MhCT08 establishment.

A recent study demonstrated the use of different mediums to enhance differential growth of epithelial and fibroblast populations (40). The cultures reported in the current study were established in RPMI-1640 complete medium so as to eliminate any phenotypic or genotypic changes arising due to patient-derived xenograft generations. The cultures did not require any feeder cells to grow and stabilized spontaneously, without the intervention of viral vectors.

Kaur et.al described an innovative method of treating the cultures with anti-fibroblast antibody along with complement rabbit serum treatment to remove fibroblast populations (32). CD90/CD44 based separation has also been successful in separating mesenchymal (CD90 positive) and epithelial (CD90 negative) populations from tumor tissue (31). Present study exploits differential trypsinization method to establish the novel autologous pair described.

PanCK and FSP-1 staining have been reported to identify epithelial and fibroblast populations exclusively (41–43). MhCT12 epithelial and fibroblast were stained with PanCK and FSP1 with high specificity. However, MhCT08, may be coming from a more aggressive tumor, probably has markers different from PanCK and FSP1 leading to lesser corresponding values of MhCT12 checked by flow cytometry.

DNA ploidy determination can correlate with the cancer grade, aggressiveness and metastatic potential of a tumor and can provide insightful information about cancer diagnosis and prognosis (44). The DNA index calculation implied the aneuploidy nature of the established cell lines, which may prove chromosomal instability, thus promoting a heterogeneous tumor evolution (45).

In vitro assays like proliferation, invasion-migration and sphere formation provide knowledge about the cancer stem cell subpopulation within a heterogeneous cell population (46,47). Both MhCT12 and MhCT08 epithelial cell lines intrinsically had sphere formation ability which was significantly increased upon treatment with corresponding fibroblast conditioned medium. This may be due to various signalling factors released by the CAFs which promote tumorigenesis and is also an example of tumor-stroma crosstalk (48,49).

The novelty of the cultures thus established was further confirmed by STR profiling. When compared to the ATCC and CLASTR databases, these cell lines did not have any significant match with the existing cell lines, thus confirming their novelty and unique origin. The fibroblast and epithelial cells showed almost similar profile confirming their autologous nature.

High risk of HPV 16 and 18 infection has been associated with oral cancer. More than 25% of all the oral cancers are associated with HPV16 infection, while a lower percentage of about 1-3% is attributed to HPV18 infection (50). Factually, HPV positive oral cancers are associated with a favourable prognosis when compared to the HPV negative oral cancers (22). Both MhCT12 and MhCT08 were shown to be HPV positive by PCR and ICC with Anti-HPV antibodie.

Since the cell lines are a pair of CAFs and cancer epithelial cells, using such model systems, specifically with co-culture using conditioned medium, will aid in understanding the molecular mechanism behind stroma-tumor interaction leading to treatment resistance, a major problem in various cancer types. The described co-culture system mimic the actual crosstalk between tumor cells and its static microenvironment, the CAFs. Moreover, the CAFs themselves, can be excellent model systems to understand tumor progression.

In summary, the cell lines developed in the present study have persistent growth potential and stable cell morphology. They hold great potential to be used as a novel drug testing platform in co-culture environment in addition to providing a useful tool to perform basic and translational research in the area of tumor-stroma interaction in human tongue cancer arising from no risk habits.

## Acknowledgements

The authors would like to thank Dr. Sujan K Dhar of MSMF for the help in analysing the STR profiling data for establishing the novelty of the established cell lines.

## Availability of data and materials

All the data generated and/or analysed is provided along with the manuscript.

## Author’s contributions

MD and AS conceived and designed the study. CG collected the patient samples and isolated fibroblast and epithelial cells. ND performed the experiments. MAK and VP provided the samples. MD and ND wrote the manuscript, analysed and interpreted the data. AS reviewed the manuscript. All authors read and approved the final manuscript.

## Ethics approval and consent to participate

The present study was approved by the ethics committee of Narayana Health City, Bangalore, Karnataka (India). The approval number is given as: NHH/MEC-CL-2015-405 (A)

## Patient consent for publication

Informed consent for the study was obtained from all the patients involved in the study.

---

**Supplementary figure 1 HPV detection.** (A) ICC based HPV infection status of primary cell lines using p16 and E7 antibodies (scale bar 50μm) (B) PCR based HPV detection in primary cell lines using MY09/MY11 primers. Beta actin is being used as the template control.

Loading order – C-: Negative No Template Control; C+: HeLa as a positive control for detection of HPV; 1: MhCT12 Epithelial; 2: MhCT12 Fibroblast; 3: MhCT08 Epithelial, and 4: MhCT08 Fibroblast

