## Supplementary figures and images for "Establishment and characterization of novel autologous pair primary cultures from two Indian non-habitual tongue carcinoma patients"

### Supplementary figure 1

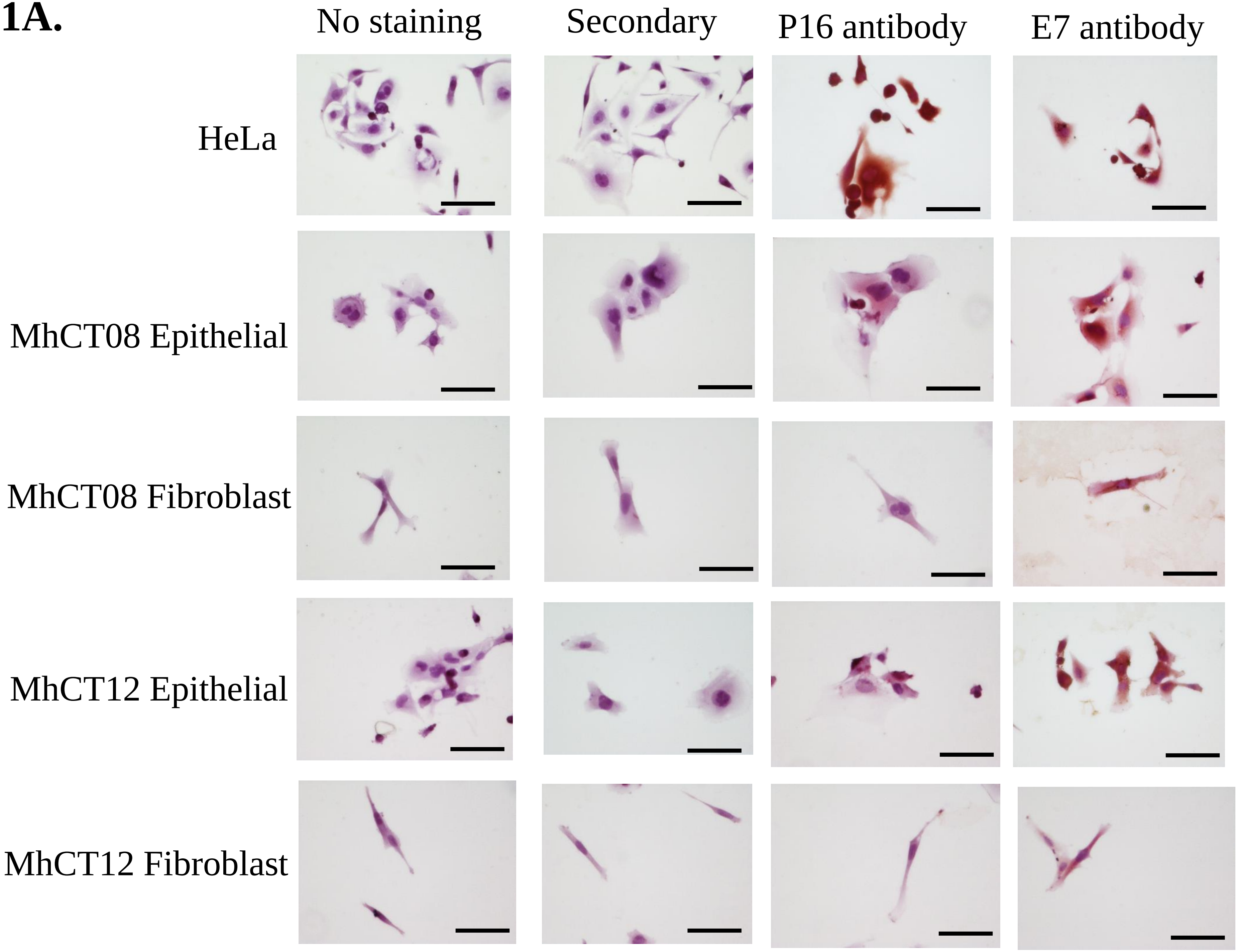

S1B.

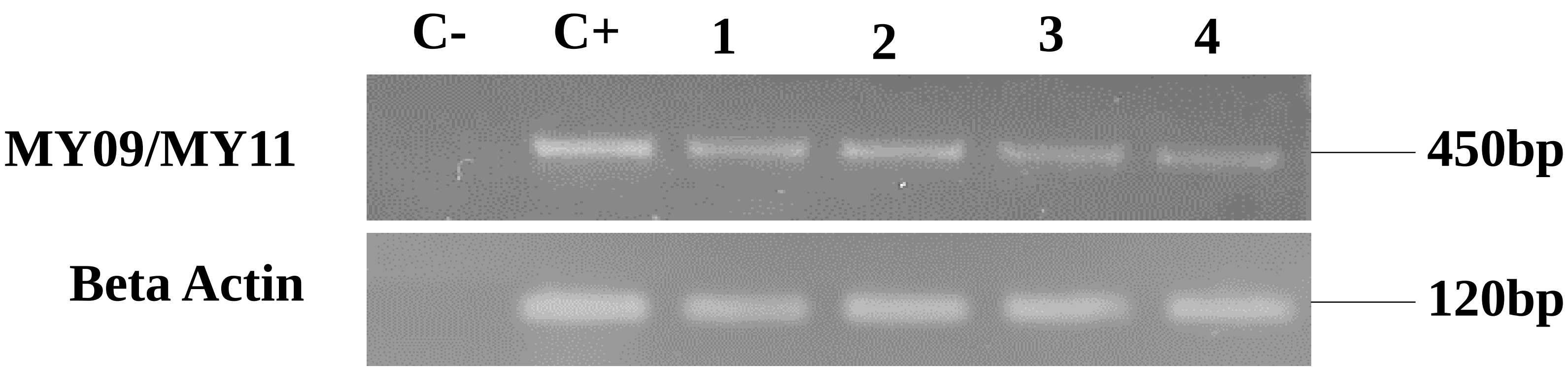
